# EFL-3/E2F7 modulates Wnt signalling through repressing the LIT-1 Nemo-like kinase during asymmetric epidermal cell division in *Caenorhabditis elegans*

**DOI:** 10.1101/2024.07.01.601538

**Authors:** Mar Ferrando-Marco, Michalis Barkoulas

## Abstract

The E2F family of transcription factors is conserved in higher eukaryotes and plays pivotal roles in controlling gene expression during the cell cycle. Most canonical E2Fs associate with members of the Dimerisation Partner (DP) family to activate or repress target genes. However, atypical repressors, such as E2F7 and E2F8, lack DP interaction domains and their functions are less understood. We report here that EFL-3, the E2F7 homologue of *C. elegans*, regulates epidermal stem cell differentiation. We show that phenotypic defects in *efl-3* mutants depend on the Nemo-like kinase *lit-1.* EFL-3 represses *lit-1* expression through direct binding to a *lit-1* intronic element. Increased LIT-1 expression in *efl-3* mutants reduces POP-1/TCF nuclear distribution, and consequently alters Wnt pathway activation. Our findings provide a mechanistic link between an atypical E2F family member and NLK during *C. elegans* asymmetric cell division, which may be conserved in other animals.

**Highlights:**

- EFL-3 is enriched in anterior daughter cells following asymmetric seam cell division.
- *efl-3* mutants show defects in cell differentiation and seam cell fate maintenance.
- EFL-3 directly represses *lit-1*/NLK expression.
- EFL-3-mediated *lit-1* repression alters POP-1 nuclear levels, linking EFL-3 to Wnt signalling.

## Introduction

Development of multicellular organisms depends on the formation of different tissues and specialised cell types. Stem cells are central in this process having the potential to give rise to cellular diversity through asymmetric cell division, while they are maintained in an undifferentiated state and can increase their population through symmetric division^1^. The balance between symmetric and asymmetric divisions needs to be tightly regulated because defects in stem cell regulation can compromise tissue homeostasis and increase the risk of cancer^2, 3^. Therefore, studying key genes and pathways regulating stem cell behaviour is central for unravelling mechanisms at play in healthy and disease state.

The epidermal seam cells of *Caenorhabditis elegans* serve as a model to study stem cell regulation^4^. Seam cells are linearly arranged throughout the length of the body (named H0-H2, V1-V6, and T) and divide symmetrically and asymmetrically in a stem-like manner during postembryonic development (Figure 1A). Following most seam cell asymmetric divisions, posterior daughter cells maintain the seam cell fate, whereas anterior daughter cells endoreduplicate and fuse to the hyp7 epidermal syncytium^5, 6^. The seam cells generate the majority of the epidermal nuclei and thus the cuticle, which is essential for growth and protection from biotic and abiotic stress^7^. The H2, V5 and T seam cell lineages also give rise to neuronal precursors^5^. The seam cells expand their population from 10 to 16 through a symmetric division at the early L2 stage, when both daughter cells maintain the proliferative potential. At the end of larval development, the 16 seam cells terminally differentiate and fuse into another syncytium that generates the cuticle ridges called alae^8^. Seam cell divisions and their position along the anterior-posterior axis are highly invariant, allowing the characterisation of phenotypic errors at single-cell resolution^9^.

**Figure 1.**
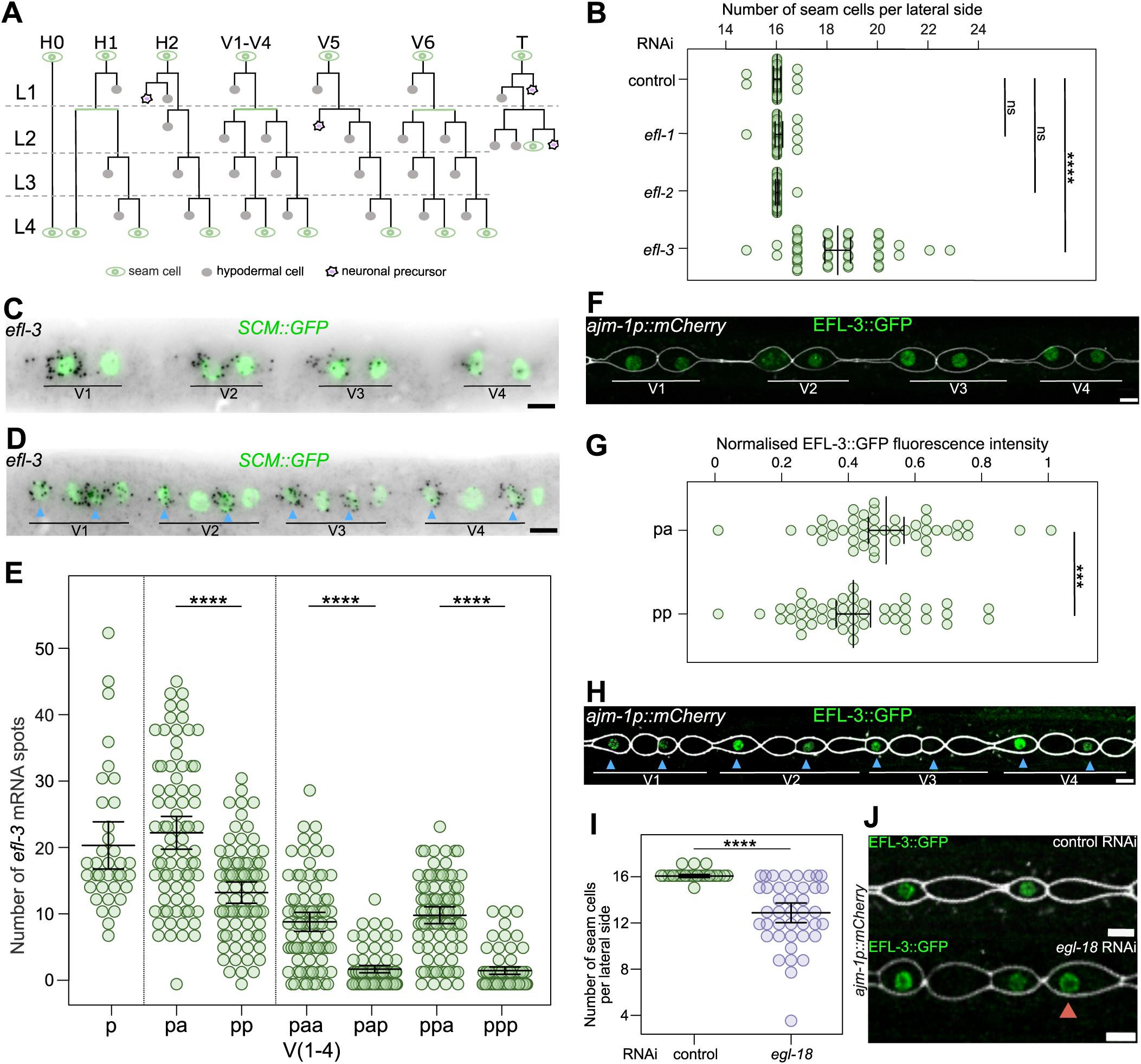
EFL-3 is an E2F transcription factor regulating seam cell development. **(A)** Schematic showing seam cell divisions from L1 to L4. Seam cells are labelled in green, hypodermal cells are labelled in grey and neuroblasts are labelled in purple. **(B)** Seam cell number counts upon RNAi targeting members of the E2F family in *C. elegans*. Unlike *efl-1* and *efl-2*, downregulation of *efl-3* leads to a significant increase in the terminal seam cell number (p<0.001 with a t-test, n = 30). **(C-D)** Representative *efl-3* smFISH images following the L2 symmetric division (C), and the L2 asymmetric division (D). Seam cell nuclei are labelled using *SCM::GFP*. **(E)** Quantification of *efl-3* mRNA spots in V1-V4 seam cells before (p) and after the L2 symmetric division (pa, pp), and following the L2 asymmetric division (paa, pap, ppa, ppp), 36 ≤ n ≤ 166. **(F)** Representative image of EFL-3::GFP before the L2 asymmetric division. **(G)** Quantification of EFL-3::GFP fluorescence intensity before the L2 asymmetric division in V1-V4 lineages. Anterior cells show higher amounts of EFL-3 compared to posterior cells (*p*<0.01 with a t-test, n = 45). **(H)** Representative image of EFL-3::GFP after the L2 asymmetric division. **(I)** *egl-18* RNAi-treated animals show a statistically significant decrease in the mean seam cell number (*p*<0.001 with a t-test) compared to control animals, n = 41. **(J)** Posterior daughter cells (marked with red arrowhead) show expression of EFL-3::GFP when treated with *egl-18* RNAi following the asymmetric L3 division, n = 91. Blue arrowheads in D and H point to anterior daughter cells. In F, H and J the membrane of the seam cells is labelled using the apical junction marker *ajm-1p::mCherry*. Error bars in B, E, G and I show the mean ± standard deviation. Scale bars are 5μm in C, D, F, H and J.

Seam cell development is controlled by Wnt signalling, a widely conserved pathway consisting of intercellular signals that can travel in the body to shape tissues through influencing cell polarity and proliferation^10–12^. Correct seam cell patterning relies on the Wnt/β-catenin asymmetry pathway (WβA). This pathway ensures that following asymmetric division, posterior daughter cells activate Wnt targets and retain seam cell fate, while anterior daughter cells do not activate Wnt and instead differentiate^13, 14^. In the dividing mother cell, Wnt ligands drive positive regulators such as Frizzled receptors and Dishevelled to localise to the posterior cortex^15–17^. Instead, negative WβA regulators, such as APR-1/APC and PRY-1/Axin asymmetrically localise to the anterior cortex^18, 19^. The β-catenin WRM-1 also localises to the anterior cortex and recruits more APR-1/APC and PRY-1/Axin^19, 20^. In turn, APR-1 promotes anterior nuclear export of WRM-1 through regulation of microtubule dynamics, while absence of APR-1 from the posterior daughter cell after division leads to nuclear WRM-1 localisation^17, 21^. Moreover, cortical APR-1 promotes SYS-1/β-catenin degradation through the destruction complex, leading to asymmetric localisation of SYS-1 to the posterior nucleus^18^ In posterior daughter cells, high nuclear WRM-1 binds to LIT-1 promoting its activation by MOM-4/MAPK and nuclear translocation^20, 22, 23^. Nuclear LIT-1 phosphorylates POP-1, the sole Tcf/Lef homologue of *C. elegans*, leading to its nuclear export^24–27^. Low POP-1 and high SYS-1 levels in posterior daughter nuclei allow the formation of a TCF/β-catenin transcriptional activator complex (POP-1/SYS-1) that activates Wnt targets^13, 28–30^, such as the GATA transcription factor EGL-18 ^31^. In anterior daughter cells, high POP-1 and low SYS-1 levels in the nucleus lead to recruitment of corepressors and transcriptional repression of Wnt targets^30, 32^. Besides Wnt signalling, seam cell patterning is also regulated by other conserved transcription factors, such as a Runx/CBFβ module, CEH-16/engrailed and various GATA factors, which can influence the decision between stem cell fate maintenance or differentiation^33–39^

The E2F family of transcription factors (E2F1-8 in humans) controls gene expression during the cell cycle. E2F1-6 associate with DP proteins to form heterodimers that control the expression of cell cycle genes^40, 41^. Atypical E2F7-8 lack a DP binding domain and are thought to act as transcriptional repressors^42–46^. Most E2Fs are regulated during the cell cycle through direct binding to pocket proteins, such as the tumour suppressor RB. These interactions are in turn regulated by the activity of cyclin-dependent kinase (CDK) and cyclin complexes, which phosphorylate pocket proteins causing E2F release^40^. The CDK-RB-E2F pathway is essential to prevent abnormal proliferation and tumour growth, with unrestrained E2F-mediated transcription being a key driver for many cancers^47, 48^. While E2Fs have been largely studied in the context of cell division, it is known that they also have non-canonical functions beyond the cell cycle ^49, 50^. The *C. elegans* genome contains three homologs of the mammalian E2F family, *efl-1*, *efl-2* and *efl-3*. EFL-1 and EFL-2 bind to DP/DPL-1 to repress expression of genes ensuring correct development of tissues such as the vulva^41, 51, 52^. For example, *efl-1* participates in the DRM complex and the SynMuvB gene regulatory pathway that inhibits vulval development and prevents somatic cells from acquiring a germline fate^53, 54^. In addition, *efl-1* and *efl-2* have been described to have pro-apoptotic function in the germline^55^.

In this study, we describe the role of EFL-3, the E2F7 homologue of *C. elegans,* in seam cell development. We report that EFL-3 localises preferentially in anterior daughter cells following asymmetric seam cell division and this localisation depends on Wnt-signalling. Tissue-specific *efl-3* mutants display gains and losses of seam cells. These phenotypes are dependent on the Nemo-like kinase *lit-1*, which we find to be a direct EFL-3 target. Increase of *lit-1* expression reduces POP-1 levels in the seam cell daughter nuclei leading to changes in seam cell fate specification. Our study therefore provides evidence that the atypical E2F EFL-3 modulates epidermal stem cell patterning in *C. elegans* through regulation of *lit-1* expression, and consequently Wnt signalling pathway activity.

## Results

### EFL-3 shows an asymmetric distribution following seam cell division

As part of our efforts to gain insights into the machinery regulating seam cell development, we previously performed transcriptional profiling and identified *efl-3* as a seam cell-expressed transcription factor^56^. The *C. elegans* genome contains three E2F family members, *efl-1*, *efl-2* and *efl-3,* but only downregulation of *efl-3* led to a significant change in the average number of seam cells (Figure 1B).

To characterise the *efl-3* expression pattern in the seam cells, we used single-molecule fluorescence *in situ* hybridisation (smFISH). We focused on the V1-V4 lineages that divide in a similar division pattern, and the L2 larval stage during which seam cells divide twice, first symmetrically and then asymmetrically. Interestingly, we observed an asymmetric localisation of *efl-3* transcripts between the two daughter cells following the symmetric and asymmetric L2 divisions (Figure 1C and D). In both cases, anterior daughter cells displayed a higher number of *efl-3* mRNA transcripts compared to posterior daughters (Figure 1E). Following the L2 asymmetric cell division, posterior daughter cells presented strong reduction in *efl-3* transcript abundance, with 50% of cells showing no *efl-3* mRNAs at all (Figure 1E).

To assess whether this asymmetric mRNA distribution also reflected how EFL-3 is distributed at the protein level, we generated and imaged an *efl-3::gfp* knock-in strain. Consistent with the mRNA transcript enrichment in anterior cells, EFL-3::GFP expression was moderately enriched in anterior daughter cells after symmetric division in comparison to posterior daughters (Figure 1F and G). Following the L2 asymmetric division, this enrichment became more pronounced, with EFL-3::GFP expression specifically observed in anterior daughter cells (Figure 1H). We reasoned that the Wnt pathway that is normally activated in posterior daughter cells retaining the seam cell fate may contribute to the *efl-3* repression seen in posterior cells. To test this hypothesis, we knocked-down the Wnt downstream effector *egl-18* by RNAi, which results in loss of seam cell fate maintenance and decreased terminal seam cell number (Figure 1I). Notably, we found EFL-3::GFP to be expressed in 21% of posterior daughter cells in *egl-18* RNAi animals compared to 0% in the control treatment (Figure 1J). Taken together, these results highlight that EFL-3 distribution in seam cells is asymmetric and that this asymmetry is dependent on Wnt pathway activation.

### Loss of *efl-3* function leads to gains and losses of seam cells during larval development

To consolidate the RNAi results, we sought to study the impact of strong loss of *efl-3* function on seam cell development. To circumvent the fact that *efl-3* is an essential gene^57^, we constructed a seam cell-specific mutant by inserting *loxP* recombination sites flanking the *efl-3* locus and introducing this modification into a genetic background expressing Cre recombinase specifically in the seam cells. We confirmed that neither the introduction of *loxP* sites flanking the *efl-3* locus nor the expression of Cre recombinase in the seam cells disrupt seam cell development (Figure S1A). Furthermore, transgenic *efl-3^loxP^* animals carrying the Cre recombinase did not show any *efl-3* expression in the seam cells by smFISH, hence they are likely to represent seam cell-specific null mutants (Figure S1B). We found that tissue-specific *efl-3* mutants displayed a stronger phenotype than *efl-3* RNAi treated animals, with both gains and losses of seam cells observed at the early adult stage (Figure 2A and B). Gains of seam cells were more frequent in the H1p and anterior V lineages (Figure 2C), whereas seam cell losses were more frequently observed in the posterior V lineages (Figure 2D). When *efl-3* mutants were treated with *egl-18* RNAi, the seam cell loss phenotype was enhanced (Figure S1C). Taken together, these results suggest that EFL-3 can both promote and repress seam cell fate.

**Figure 2.**
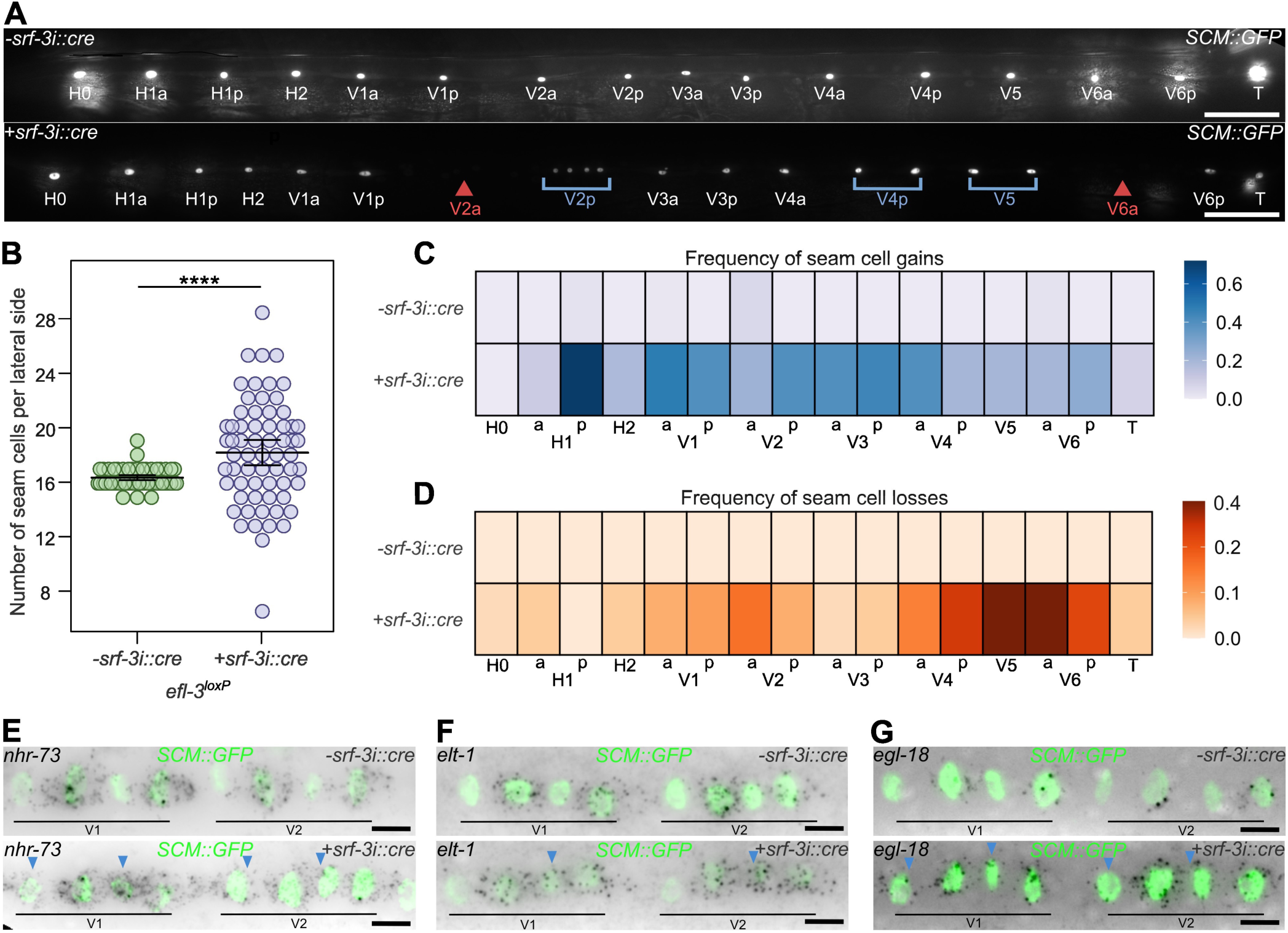
A tissue-specific *efl-3* mutant displays gains and losses of seam cells. **(A)** Representative images of tissue-specific *efl-3* mutant animals (+*srf-3i::cre efl-3^loxP^*) versus controls (*-srf-3i::cre efl-3^loxP^*) at the early adult stage. Blue brackets indicate lineages with seam cell gains, red arrowheads point to lineages with seam cell losses. **(B)** Seam cell counts in tissue-specific *efl-3* mutant animals show a significant increase in the average number of seam cells in comparison to controls (p<0.001 with a t-test, 58 ≤ n ≤ 64). Error bars show the mean ± standard deviation. **(C-D)** Heatmaps showing frequency of seam cell gains (C) and losses (D) per cell lineage at the end of postembryonic development in tissue-specific *efl-3* mutant animals versus controls (n = 50 per condition, a = anterior lineage, p = posterior lineage). **(E-G)** Representative smFISH images of *nhr-73* (E), *elt-1* (F) and *egl-18* (G) mRNA distribution in tissue-specific *efl-3* mutant animals in comparison to controls following the L2 asymmetric division. Blue arrowheads point to anterior daughter cells that ectopically express *nhr-73*, *elt-1* and *egl-18* in *efl-3* mutant animals. Seam cell nuclei are labelled using *SCM::GFP*. Scale bars are 50 μm in A and 5μm in E-G. See also Figure S1.

To determine at what larval stage seam cells are gained or lost in the *efl-3* mutant, we carried out detailed lineage analysis during post-embryonic development. We found that at the late L2 and late L3 stages, *efl-3* mutant animals only displayed seam cell gains (Figure S1D, E and F). Gaps in the seam cell line were only observed later at the late L4 stage, as shown by the loss of expression of the epidermal marker *wrt-2p::GFP-PH* (Figure S1G). Cells within these gaps also lacked expression of another seam cell marker, the nuclear hormone receptor *nhr-73* by smFISH (Figure S1H). Therefore, lineage analysis suggests that in *efl-3* mutant animals seam cell losses occur only following the L4 asymmetric division and thus later than the onset of gains.

To characterise the developmental state of the anterior V1-V4 cells that retain the expression of *SCM::GFP* following division, we investigated the expression pattern of other transcription factors whose expression is known to correlate with seam cell fate, such as *nhr-73*, *elt-1* and *egl-18.* In all cases, we found that *efl-3* mutants ectopically expressed these genes in anterior daughter cells following asymmetric division (Figure 2E-G). In the case of *egl-18*, we confirmed this result at the protein level by studying the expression of an endogenously tagged *egl-18* with *mNeonGreen*. We found a significant increase in EGL-18 protein expression in anterior daughter cells in *efl-3* mutants compared to control (Figure S1I and J).

Because E2Fs, including E2F7, are known to participate in the regulation of the cell cycle ^58, 59^, we asked whether the defects in cell differentiation in *efl-3* mutant animals may be an indirect consequence of cell cycle perturbation. Lineage analysis did not reveal any gross difference in the timing of cell division between wild-type and mutant. Furthermore, it is known that following the L2 asymmetric division, anterior daughter cells endoreduplicate before fusing to the hyp7 syncytium^8^. We found that *efl-3* mutants did not show changes in endoreduplication in anterior nuclei (Figure S1K and L). Taken together, these findings suggest that the observed phenotypes in *efl-3* mutants are more likely to be linked directly to changes in cell differentiation.

### Identification of putative EFL-3 targets via NanoDam

To identify downstream targets of EFL-3 in the seam cells, we used a modified TaDa approach for streamlined tissue-specific transcription factor binding called NanoDam^60, 61^. NanoDam employs the main TaDa principle to overcome the considerable toxicity observed when Dam-fusions are expressed at high levels in animal cells^35, 62^. Briefly, transgenes are expressed at very low levels in the tissue of interest via a bicistronic mRNA cassette design harbouring two open reading frames (ORFs). The first ORF encodes a fluorescent protein and is followed by two STOP codons and a frameshift mutation before the ATG of the second ORF. Because of the universal property of eukaryotic ribosomes to reinitiate translation after a STOP codon at a reduced frequency, this transgene configuration leads to low levels of Dam-Protein of interest fusion production. NanoDam relies on the tissue-specific expression of a vhhGFP4 nanobody fused to Dam. This allows profiling of putative targets at the desired tissue upon crossing the GFP nanobody:Dam transgene to a TF-GFP transgenic line (here the *efl-3::gfp* knock-in strain). We performed EFL-3 NanoDam at the L4 stage using both C-terminal and N-terminal fusions of the Dam:GFP-nanobody transgene expressed in the seam cells. The functionality of the EFL-3::GFP fusion was assessed by scoring the seam cell number in heterozygous +*srf-3i::cre efl-3^loxP^* / *efl-3::gfp* animals, which showed a wild-type seam cell number similar to heterozygous +*srf-3i::cre efl-3^loxP^/ wt* animals (Figure S2A)

Amplicons representing methylated genomic sequences were sequenced and data was analysed using the damidseq pipeline^63^. EFL-3 NanoDam signal was calculated as log2 (EFL-3:GFP; Dam:GFP-nanobody / Dam:GFP-nanobody) ratio scores per GATC fragment of the genome for each Dam:GFP-nanobody configuration. Aggregate genome-wide signal profiles across genes containing statistically significant peaks between 5 kb upstream of the transcriptional start site (TSS) and 2 kb downstream of the transcription end site (TES) showed signal enrichment in sequences proximal to the TSS and the TES (Figure S2B). We identified 1316 and 1237 statistically significant peaks in the case of the C- and N-terminal Dam:GFP-nanobody fusions, respectively (Table S1), with > 70% of peak overlap between the two datasets (Figure S2C). Hierarchical clustering of the localization and score of those peaks that lie between 5 kb upstream and 2 kb downstream of genes revealed clusters of signal enrichment upstream to TSS, within the genes and downstream to TES regions (Figure S2D). Peaks were assigned to the closest gene leading to 1882 and 1826 genes for each configuration with an 80% overlap between the two gene datasets. The list of candidate EFL-3 target genes showed significant overlap with genes expressed in seam cells based on sci-RNAseq data^64^ and with a filtered list of genes with known roles in seam cell development from the literature (Table S2).

### EFL-3 controls *lit-1* expression in seam cells

One potential target reproducibly identified in the NanoDam datasets was *lit-1*, which encodes a MAP kinase homolog of the human NLK and is known to participate in Wnt signal transduction by regulating the nuclear distribution of POP-1^20, 24–27^. NanoDam profiling revealed significant signal enrichment in the first intron of *lit-1* (Figure 3A). We carried out *lit-1* smFISH to compare *lit-1* expression between wild type and *efl-3* mutants after the L2 asymmetric division. We found a significant increase in *lit-1* transcript levels in anterior and posterior daughter cells, suggesting that EFL-3 may act as a repressor of *lit-1* expression in the seam cells (Figure 3B). Following an asymmetric seam cell division, LIT-1 has been reported to localise preferentially to the nucleus of posterior daughter cells where it phosphorylates POP-1 to trigger its nuclear export^20^. We therefore investigated LIT-1 nuclear levels during symmetric and asymmetric L2 division using a *lit-1::gfp* knock-in strain. Consistent with the smFISH results above, we found that LIT-1 levels were significantly increased in *efl-3* mutant animals in anterior and posterior daughter cells following symmetric and asymmetric division (Figure 3C and D, Figure S3A and B).

**Figure 3.**
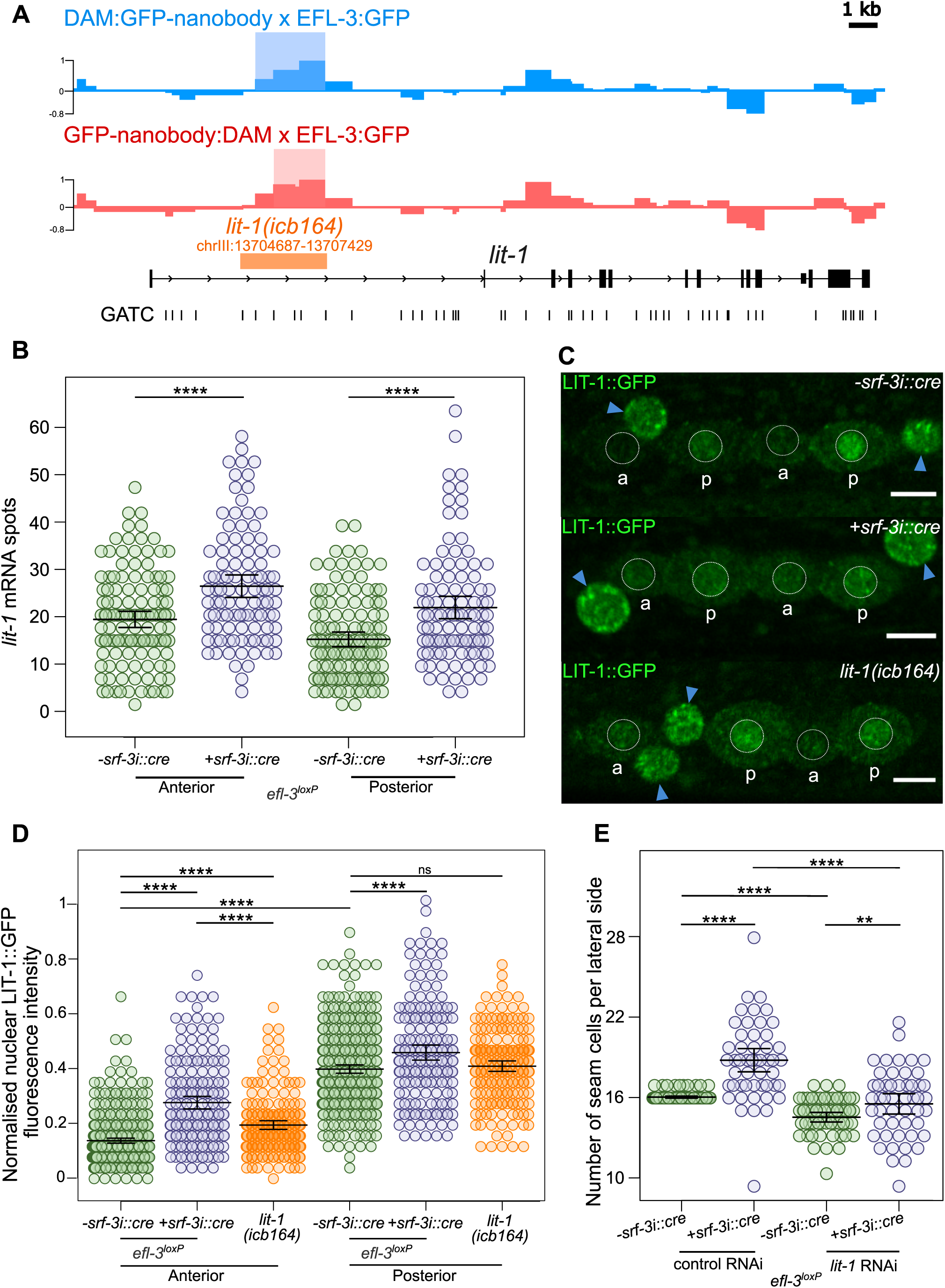
EFL-3 directly represses *lit-1* expression. **(A)** Signal profiles for C-terminal (blue) and N-terminal (red) NanoDam reflecting EFL-3::GFP binding near the *lit-1* locus. The y axis represents log_2_(EFL-3::GFP; GFP-nanobody:dam /GFP-nanobody:dam) score and the scale bar length is 1 kb. Statistically significant peaks are indicated by lightly shaded rectangles on the signal tracks (FDR < 0.05). The generated *lit-1(icb164)* CRISPR deletion is shown in orange. **(B)** Quantification of *lit-1* mRNA spots in V1-V4 lineages in anterior and posterior daughter cells in *efl-3* tissue-specific mutants (*+srf-3i::cre efl-3^loxP^*) versus controls (*-srf-3i::cre efl-3^loxP^*) following the L2 asymmetric division, 100≤ n ≤ 126 per condition. **(C)** Representative images of LIT-1::GFP following the L2 asymmetric division in controls (*-srf-3i::cre efl-3^loxP^*), tissue-specific *efl-3* mutants (*+srf-3i::cre efl-3^loxP^*) and animals containing the *lit-1(icb164)* CRISPR deletion. Seam cell nuclei are circled in white. Hypodermal nuclei are labelled with blue arrowheads, a = anterior daughter cell, p = posterior daughter cell. **(D)** Quantification of LIT-1::GFP fluorescence intensity in the nuclei of V1-V4 seam cells following the L2 asymmetric division in controls (*-srf-3i::cre efl-3^loxP^*), tissue-specific *efl-3* mutants (*+srf-3i::cre efl-3^loxP^*) and animals containing the *lit-1(icb164)* CRISPR deletion; n ≥ 175. **(E)** Seam cell counts in *efl-3* tissue-specific mutant animals versus controls treated with *lit-1* RNAi or empty vector; n = 50 per condition. Error bars in B, D and E show the mean ± standard deviation and **** represent *p*<0.001 and ** *p*<0.01 with a t-test.. Scale bars are 5 μm in C. See also Figures S2 and S3.

To address whether the increase in *lit-1* levels could be causative for the seam cell phenotype observed in *efl-3* mutants, we carried out *lit-1* RNAi in the *efl-3* mutant background. We found a significant reduction in the mean seam cell number of *efl-3* mutant animals upon *lit-1* RNAi treatment compared to the control treatment (Figure 3E). Detailed lineage analysis revealed that this reduction in seam cell number was driven by a statistically significant suppression of seam cell gains in anterior V lineages in *efl-3* mutants (Figure S3C). Furthermore, we found a statistically significant reduction in posterior V lineage cell losses in *efl-3* mutants treated with *lit-1* RNAi (Figure S3D), which was masked when looking at the aggregate seam cell number counts. The suppression of seam cell gains and losses observed in *efl-3* mutants suggests that both phenotypes are LIT-1-dependent.

To test whether EFL-3 could directly control the expression of *lit-1* through the identified site on the first intron, we used CRISPR-mediated genome editing to delete the entire EFL-3 NanoDam peak (Figure 3A). Animals containing this intronic deletion showed an increase in nuclear LIT-1 levels in anterior daughter cells following the asymmetric L2 division (Figure 3C and D). We note that this increase was lesser in magnitude than what is observed in *efl-3* mutants, suggesting that this intronic element may not be the only way via which EFL-3 controls *lit-1* expression in the seam cells. Consistent with this hypothesis, *efl-3* RNAi was found to further increase LIT-1 levels in animals carrying the intronic deletion. This further suggests that EFL-3 is likely to regulate *lit-1* expression directly through additional regulatory elements or indirectly through other intermediate transcription factors (Figure S3E and F).

### Decrease in nuclear POP-1 levels correlates with seam cell gains and losses in *efl-3* mutants

LIT-1 accumulation in the nucleus of posterior daughter cells is predicted to cause POP-1 nuclear export, thereby ensuring an optimal β-catenin (SYS-1) / POP-1 ratio in the nucleus that is conductive to Wnt target activation^20, 28–30^. Low LIT-1 levels in the nuclei of anterior daughter cells allow POP-1 to also remain at high levels in the nucleus and associate with proteins repressing Wnt target genes^30, 32^. We therefore asked whether higher LIT-1 levels in *efl-3* mutant nuclei have a consequence on POP-1 localisation. To test this prediction, we studied nuclear POP-1 protein levels upon loss of *efl-3* function using a *gfp::pop-1* knock-in strain. In wild-type animals, POP-1 levels were lower in posterior daughter cells in comparison to anterior daughter cells following the L2 asymmetric division (Figure 4A and B). In *efl-3* mutant animals, we found a significant decrease in POP-1 levels in both anterior and posterior daughter cells (Figure 4A and B). Changes in POP-1 expression were at the protein level because *pop-1* mRNA transcripts were unchanged between control and *efl-3* RNAi-treated animals (Figure S4A and B). We also compared POP-1 nuclear levels at the L2 and L4 stages. In wild type animals, POP-1::GFP expression was higher at the L4 stage compared to L2 (Figure S4C and D). Additionally, we observed a less pronounced asymmetry between anterior and posterior daughter cells at the L4 stage. (Figure S4E). POP-1 levels were reduced to a similar average level in *efl-3* mutants at both the L2 and L4 stages (Figure S4 C and D). These results suggest that while reduction in POP-1 levels can readily explain the seam cell gains observed in *efl-3* mutants, absolute POP-1 levels alone cannot explain the seam cell losses observed specifically at L4.

**Figure 4.**
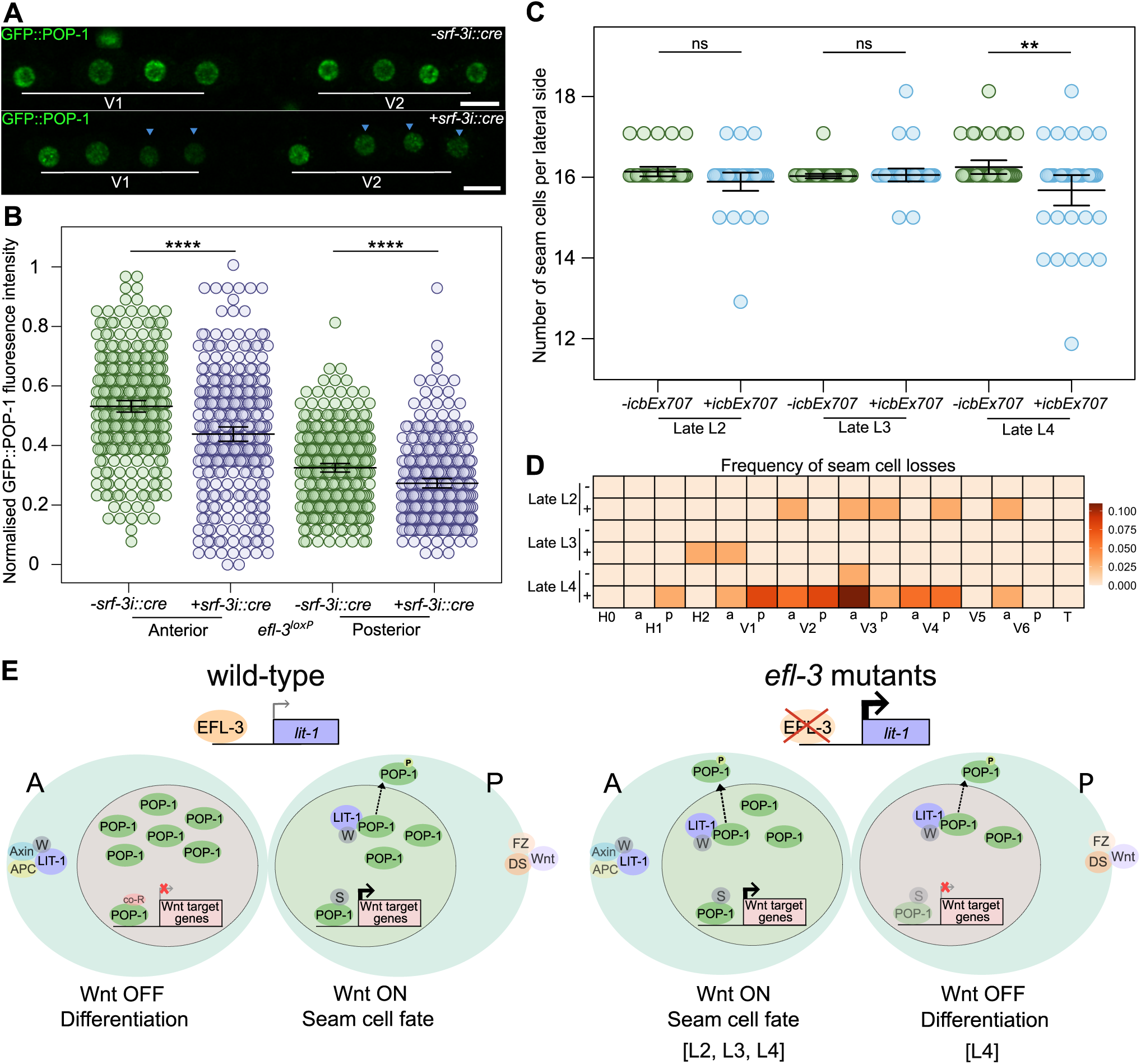
*efl-3* mutants show changes in POP-1 nuclear distribution that are compatible with seam cell gains and losses. **(A)** Representative images of GFP::POP-1 following the L2 asymmetric division in tissue-specific *efl-3* mutant animals (*+srf-3i::cre efl-3^loxP^*) versus controls (*-srf-3i::cre efl-3^loxP^*). Blue arrowheads indicate cells with visually lower expression of GFP::POP-1. Scale bars are 5 μm. **(B)** Quantification of GFP::POP-1 fluorescence intensity following the L2 asymmetric division in tissue-specific *efl-3* mutant animals versus controls, n ≥ 294. **** *p*<0.001 with a t-test. **(C)** Seam cell counts in wild-type (-*icbEx707*) and animals overexpressing *lit-1* under the *arf-5i* element (+*icbEx707*) at the late L2, late L3 and late L4 stages. At late L4, +*icbEx707* present a significant decrease in the average number of seam cells compared to wild-type (\*\**p*<0.01 with a t-test), n = 36. **(D)** Heatmap showing frequency of seam cell losses per cell lineage at the late L2, late L3 and late L4 stages in wild-type (-*icbEx707*) and in animals overexpressing *lit-1* (+*icbEx707*), n = 36 per condition, a = anterior lineage, p = posterior lineage. **(E)** Model of Wnt signalling activation in wild-type and in *efl-3* mutant animals. In wild-type animals, anterior daughters present high POP-1 nuclear levels leading to repression of Wnt target genes and differentiation. Posterior daughter cells exhibit POP-1 nuclear export thanks to LIT-1 and WRM-1 activities. POP-1 interacts with SYS-1 to drive expression of Wnt target genes resulting in seam cell fate maintenance. In *efl-3* mutants, nuclear LIT-1 levels increase in both daughter cells leading to increased POP-1 nuclear export. As a consequence, in L2, L3 and L4 divisions anterior daughter cells show ectopic activation of the Wnt signalling causing gains of seam cells. Because of additional sensitisation, in L4 divisions some posterior daughter cells fail to activate the Wnt signalling and lose seam cell fate. Fz = Frizzled; DS = Dishevelled; W = WRM-1; S = SYS-1; co-R = POP-1 co-repressors. Frizzled and dishevelled localise to the posterior cortex where they bind the Wnt ligand (Wnt). APC (APR-1), Axin (PRY-1), WRM-1 and LIT-1 localise to the anterior cortex. Error bars in B and C show the mean ± standard deviation. See also Figure S4.

To test whether increase in *lit-1* expression can explain the seam cell losses observed in *efl-3* mutants, we overexpressed *lit-1* under the *arf-5i* element, which predominantly drives expression in posterior daughters retaining seam cell fate^65^. Animals overexpressing *lit-1* presented losses of seam cells leading to a significant decrease in the mean seam cell number (Figure 4C). Similar to *efl-3* mutants, losses of seam cells were predominantly observed after the L4 asymmetric division (Figure 4C and D). As we found comparable L2 and L4 POP-1 levels in posterior daughter cells of *efl-3* mutants, we hypothesised that the losses observed at L4 may reflect an increased sensitivity to change. To test this hypothesis further, we investigated seam cell defects in a strain carrying the hypomorphic *pop-1(hu9)* allele and observed gains of seam cells leading to a significant increase in the mean seam cell number (Figure S4F). Seam cell gains were only observed at the L4 stage (Figure S4G), suggesting that seam cells are likely to present a higher sensitivity to both gains and losses at this late larval stage.

## Discussion

The E2F family of transcription factors are important for many cellular processes, including cell cycle regulation, apoptosis, and DNA damage^50, 66–68^. Atypical E2Fs are known to act as transcriptional repressors antagonising canonical E2F activators^69–71^. For example, E2F7 has been implicated in restraining cell proliferation in human cells via directly regulating the expression of genes that regulate mitotic progression among other targets, such as microRNAs^45, 58, 72, 73^. Previous work on *efl-3* in *C. elegans* suggested a role in preventing apoptosis in PVA and PVB neurons of the ventral nerve cord by repressing the expression of the pro-apoptotic gene *egl-1*^57^. In addition, *efl-3* has been implicated in the development of the somatic gonad by specifying the correct amount of distant tip cells^74^. However, the exact molecular mechanisms of how this is achieved still need to be elucidated. We report here that EFL-3 regulates asymmetric divisions in the *C. elegans* epidermal stem cell model. Consistent with a role as a transcriptional repressor, we show that EFL-3 represses *lit-1* expression in seam cells partially through direct binding to an intronic element, which has a knock-on effect on POP-1 distribution. Our study therefore reveals a link between an atypical E2F repressor and NLK-mediated regulation of Wnt signalling in the context of asymmetric cell division.

We characterised the *efl-3* expression pattern at the mRNA and protein level and report an enrichment in anterior seam cell daughters after symmetric and asymmetric divisions. This enrichment profile is consistent with a role for EFL-3 in facilitating cell differentiation of anterior daughter cells. While this function may be key to asymmetric cell division, it could be bypassed during symmetric division similar to how Wnt asymmetry is also masked at the phenotypic level^75^. Expression of atypical E2Fs is known to be cell cycle-dependent, often with a maximum expression at the S phase^45, 76^ Following seam cell asymmetric division, anterior daughters immediately proceed to S phase before fusing to the hyp7 syncytium and differentiate, whereas the cell cycle progresses more slowly in posterior daughter cells that will continue to divide^77^ This difference in cell cycle commitment between the two daughter cells could therefore contribute to the higher *efl-3* expression found in anterior daughter cells after asymmetric division. However, it is unlikely to be the main driver because a similar anterior *efl-3* enrichment was observed following the symmetric division when both daughter cells progress simultaneously to S phase^77^ We propose that the anterior enrichment is likely driven by Wnt-dependent repression in posterior daughter cells following an asymmetric cell division. This is reminiscent of the atypical E2F7 repression by Wnt signalling reported in proliferating hepatocytes^78^. Similar to *efl-3* mRNA distribution, *pop-1* transcription was also found to be stronger in anterior daughter cells post-division, which may also reflect negative feedback from Wnt activation on *pop-1* expression. Our findings indicate that besides the known regulation of the LIT-1 - POP-1 axis at the protein level^22, 23^, transcriptional regulation further safeguards seam cell fate decisions.

NLK-mediated Tcf/Lef phosphorylation can have positive or negative consequences on its activity depending on the cellular context^79–81^. In *C. elegans*, LIT-1 promotes POP-1 nuclear export allowing low nuclear POP-1 levels in posterior daughter cells and Wnt signalling activation^24–27^. In *efl-3* mutants, we found higher levels of nuclear LIT-1 and a reduction in nuclear POP-1 levels in both anterior and posterior daughter cells following asymmetric division. LIT-1 activity is regulated by WRM-1 and MOM-4. WRM-1 promotes LIT-1 autophosphorylation, facilitates phosphorylation by MOM-4 and it’s essential for LIT-1 translocation to the nucleus^22, 23^. This suggests that MOM-4 and WRM-1 activities are sufficient to handle the increased amounts of LIT-1 in *efl-3* mutants. A decrease in POP-1 levels has been shown to lead to ectopic Wnt signalling activation in anterior daughter cells and subsequent symmetrisation of seam cell divisions^11^. Following asymmetric division, posterior daughter cells require low levels of nuclear POP-1 to maintain the seam cell fate. However, if levels of POP-1 drop below a threshold, seam cell fate can no longer be maintained^75^. Therefore, changes in POP-1 nuclear levels through increased *lit-1* expression may contribute to both the seam cell gains and losses observed in *efl-3* mutants. (Figure 4E). In this scenario, lowering POP-1 levels in anterior daughter cells where levels are normally high can activate Wnt targets and prevent cell differentiation. Meanwhile, lowering POP-1 in posterior daughter cells that normally experience low POP-1 levels can lead to premature differentiation (Figure 4E). While changes in POP-1 levels could contribute to both seam cell gains and losses, our results demonstrate that absolute POP-1 levels alone cannot fully explain the *efl-3* mutant phenotypes, particularly the loss of seam cells at the L4 stage. These losses may rely upon additional sensitisation of seam cells to changes in POP-1 levels at this developmental stage. Such sensitisation may stem from changes in the abundance or distribution of Wnt signalling components between the daughter cells as animals progress through development. For example, the negative regulator PRY-1 has been reported to be more widely distributed in seam cells at the L4 stage in comparison to earlier developmental stages^82^. Alternatively, sensitisation to changes in POP-1 levels may also depend on interactions between spatial patterning machinery in seam cells and the heterochronic pathway defining temporal identities^83, 84^.

E2F-mediated regulation of proliferation and Wnt signalling are fundamental pathways in healthy development with important implications in disease such as cancer^47, 48, 85^. However, the exact crosstalk between E2Fs and Wnt signalling is not well understood. One study recently reported that atypical E2F8 knockdown inhibits the Wnt signalling pathway in ovarian cancer cells^86^. Canonical E2F1 is also known to activate expression of axin2, inhibiting Wnt signalling to promote cell death in mammalian cells ^87^. E2F1 has been shown to activate the expression of the Wnt receptor Frizzled in human osteoblasts to facilitate cell differentiation^88^. Our findings add to these reports by providing a mechanistic link between EFL3/E2F7 and the NLK-Tcf/Lef pathway in the context of epidermal stem cell fate determination in *C. elegans*. Besides TCF, NLK phosphorylates many other substrates so a link between E2F7 and NLK may have broader implications for vertebrate and invertebrate development.

## Supplementary Material

**Figure S1:**
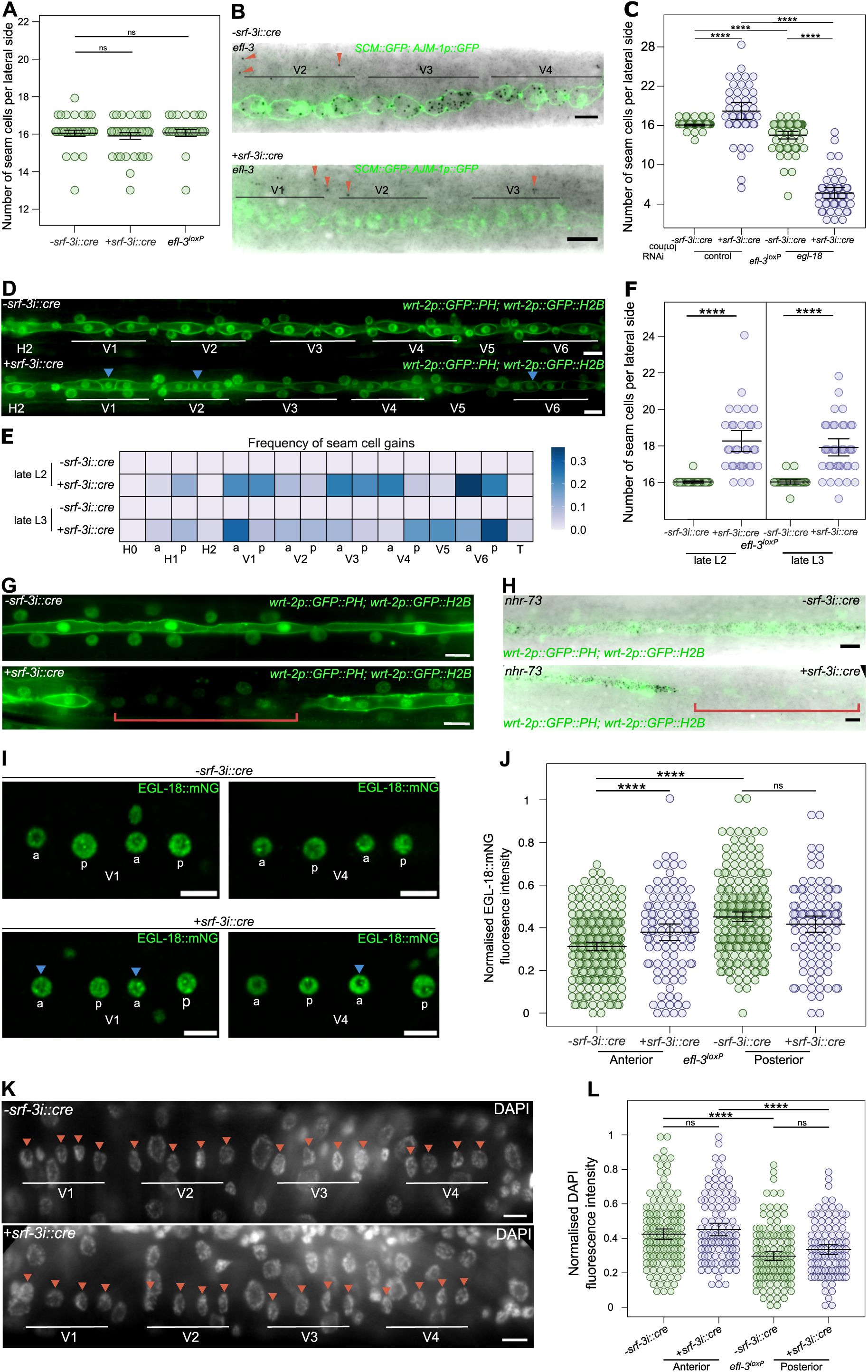
Phenotypic characterisation of *efl-3* mutants. **(A)** Seam cell counts in wild-type *(-srf-3::cre*), animals expressing the CRE recombinase in the seam cells (*+srf-3::cre*) and animals with *efl-3* locus floxed, n ≥ 30. **(B)** Representative images of *efl-3* smFISH following the L2 asymmetric division in wild-type (*-srf-3::cre efl-3^loxP^*) and in *efl-3* mutant animals (*+srf-3::cre efl-3^loxP^*). Seam cell nuclei are labelled using *SCM::GFP* and the apical junction marker *AJM-1p::GFP. +srf-3::cre* animals don’t show *efl-3* mRNA spots in the seam cells. Red arrowheads point to *efl-3* mRNA spots in the hypodermis. **(C)** Seam cell counts in wild-type and *efl-3* mutant animals treated with control and *egl-18* RNAi. Error bars show the mean ±standard deviation, n ≥ 46 per condition. **(D)** Representative images of seam cells at the late L2 stage in wild type and in *efl-3* mutant animals. Seam cells are labelled with the epidermal marker *wrt-2p::GFP::PH; wrt-2p::GFP::H2B*. Blue arrowheads indicate anterior daughter cells that failed to differentiate following asymmetric division. **(E)** Heatmaps showing frequency of seam cell gains per cell lineage at the late L2 and late L3 stages in wild-type and in *efl-3* mutant animals. No seam cell losses were observed in any of these stages, n ≥ 30 per condition, a = anterior lineage, p = posterior lineage. **(F)** Seam cell counts in wild-type and *efl-3* mutant animals at the late L2 and late L3 stages. *efl-3* mutants present a significant increase in the average number of seam cells compared to wild-type. **(G)** Representative images of the seam cells at L4 stage in wild-type (*-srf-3::cre efl-3^loxP^*) and in *efl-3* mutant animals (*+srf-3::cre efl-3^loxP^*). Seam cells are labelled with the epidermal marker *wrt-2p::GFP::PH; wrt-2p::GFP::H2B*. **(H)** Representative *nhr-73* smFISH images at L4 stage in wild-type (*-srf-3::cre efl-3^loxP^*) and in *efl-3* mutant animals (*+srf-3::cre efl-3^loxP^*). In G and H, *efl-3* mutants present gaps in the seam cell line indicated with a red bracket. **(I)** Representative images of EGL-18::mNeonGreen following the L2 asymmetric division in wild-type (*-srf-3::cre efl-3^loxP^*) and in *efl-3* mutant animals (*+srf-3::cre efl-3^loxP^*). Blue arrowheads indicate representative anterior daughter cells with higher expression of EGL-18::mNeonGreen. **(J)** Quantification of EGL-18::mNeonGreen following the L2 asymmetric division in wild-type (*-srf-3::cre efl-3^loxP^*) and in *efl-3* mutant animals (*+srf-3::cre efl-3^loxP^*), n ≥ 100. **(K)** Representative images of DAPI staining following L2 asymmetric division in wild-type (*-srf-3::cre efl-3^loxP^*) and in *efl-3* mutant animals (*+srf-3::cre efl-3^loxP^*). Seam cell nuclei are labelled with red arrowheads. **(L)** Quantification of DAPI fluorescence intensity in anterior and posterior daughter cells following L2 asymmetric division in wild-type and *efl-3* mutant animals, n ≥ 66. Error bars in A, C, F, J and L show the mean ±standard deviation. Scale bars are 5 μm in B, D, G, H, I and K. In C, F, J and L, **** *p*<0.001 with a t-test.

**Figure S2:**
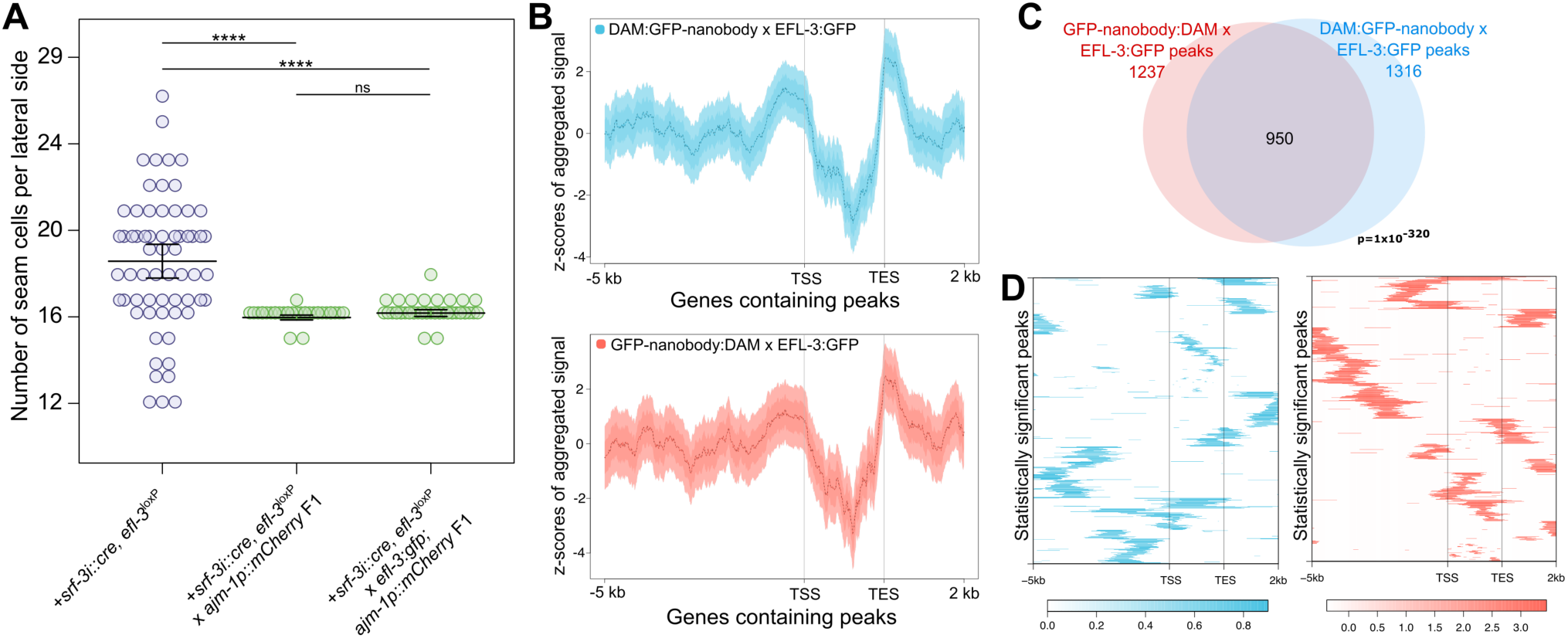
NanoDam analysis of EFL-3::GFP binding. **(A)** Seam cell counts in *efl-3* mutants, and in the F1 progeny of *efl-3* mutants crossed with WT or with *efl-3::gfp* animals. Rescue of the phenotype demonstrates the functionality of the EFL-3::GFP fusion. Cross progeny was identified via expression of the apical junction marker *ajm-1p::mCherry*, **** p<0.001 with a t-test. **(B)** Aggregation plots of profiling data generated by EFL-3 NanoDam showing average enrichment scores in 10-bp bins for regions of equal length across all genes containing statistically significant peaks. Enrichment is seen upstream and downstream of gene regions. Plots show 5 kb upstream of the transcription start site (TSS) and 2 kb downstream of the transcription end site (TES). Gene bodies are condensed into a 2 kb pseudo-length. Shaded areas represent 95% confidence intervals. **(C)** Venn diagram of all call peaks between EFL-3 C- and N-terminal NanoDam. P value was calculated by Monte Carlo simulations for significance of overlap between datasets. **(D)** Heatmaps representing the hierarchically clustered localization and enrichment score of all statistically significant peaks (FDR < 0.05) within 5 kb upstream and 2 kb downstream of genes containing peaks.

**Figure S3:**
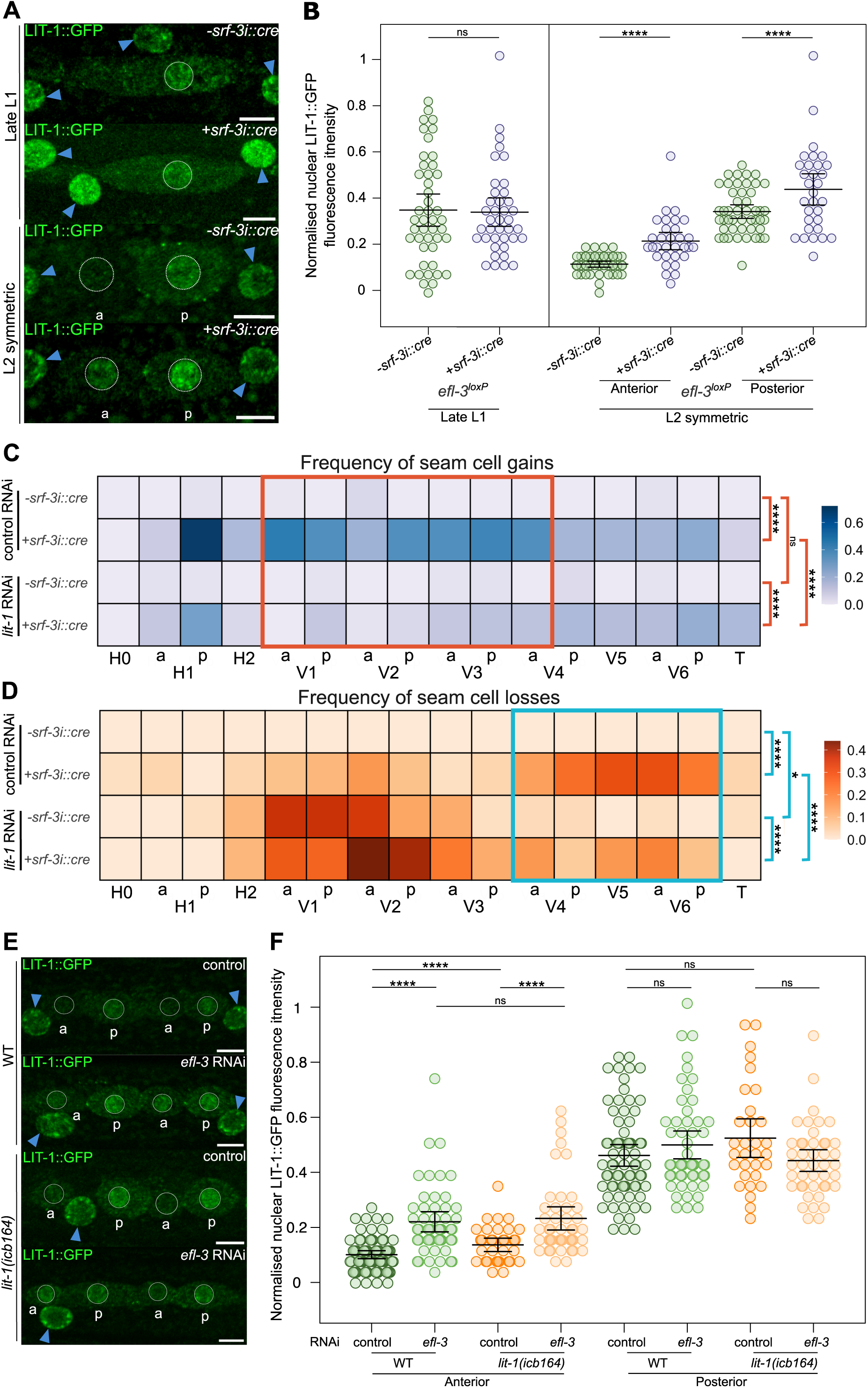
EFL-3 regulates LIT-1 levels and *lit-1* downregulation reduces the gains and losses of seam cells observed in *efl-3* mutants. **(A)** Representative images of LIT-1::GFP at late L1 stage and following the L2 symmetric division in wild-type (*-srf-3::cre efl-3^loxP^*) and *efl-3* mutant animals (*+srf-3::cre efl-3^loxP^*) **(B)** Quantification of LIT-1::GFP fluorescence intensity in the nuclei of V1-V4 seam cells at late L1 and after the L2 symmetric division. LIT-1::GFP is significantly increased in anterior and posterior daughter cells in *efl-3* mutant animals (*+srf-3::cre efl-3^loxP^*) compared to wild-type animals (*-srf-3::cre efl-3^loxP^*) after the L2 symmetric division, 31 ≤ n ≤ 47. **(C-D)** Heatmaps showing frequency of seam cell gains **(C)** and losses **(D)** per cell lineage at the end of postembryonic development in wild-type (*-srf-3::cre efl-3^loxP^*) and in *efl-3* mutant animals (*+srf-3::cre efl-3^loxP^*) treated with control and *lit-1* RNAi. Chi-square test was performed for the pooled cell lineages highlighted in the red square in C and blue square in D. * p<0.05, **** p<0.001. In C and D, n = 50 per condition. **(E)** Representative images of LIT-1::GFP following the L2 asymmetric division in wild-type and *lit-1(icb164)* animals treated with control and *efl-3* RNAi. (F) Quantification of LIT-1::GFP fluorescence intensity in the nuclei of V1-V4 seam cells following the L2 asymmetric division. LIT-1::GFP is significantly increased in anterior daughter cells in *efl-3* RNAi-treated animals compared to control animals. Animals containing *lit-1(icb164)* CRISPR deletion didn’t show further increase in LIT-1::GFP fluorescence intensity in anterior daughter cells compared to wild-type when treated with *efl-3* RNAi, 35 ≤ n ≤ 74. In A and E, seam cell nuclei are circled in white, hypodermal nuclei are labelled with blue arrowheads, a = anterior daughter cell and p = posterior daughter cell and scale bars are 5 μm. Error bars in B and F show the mean ±standard deviation. **** p<0.001 with a t-test.

**Figure S4:**
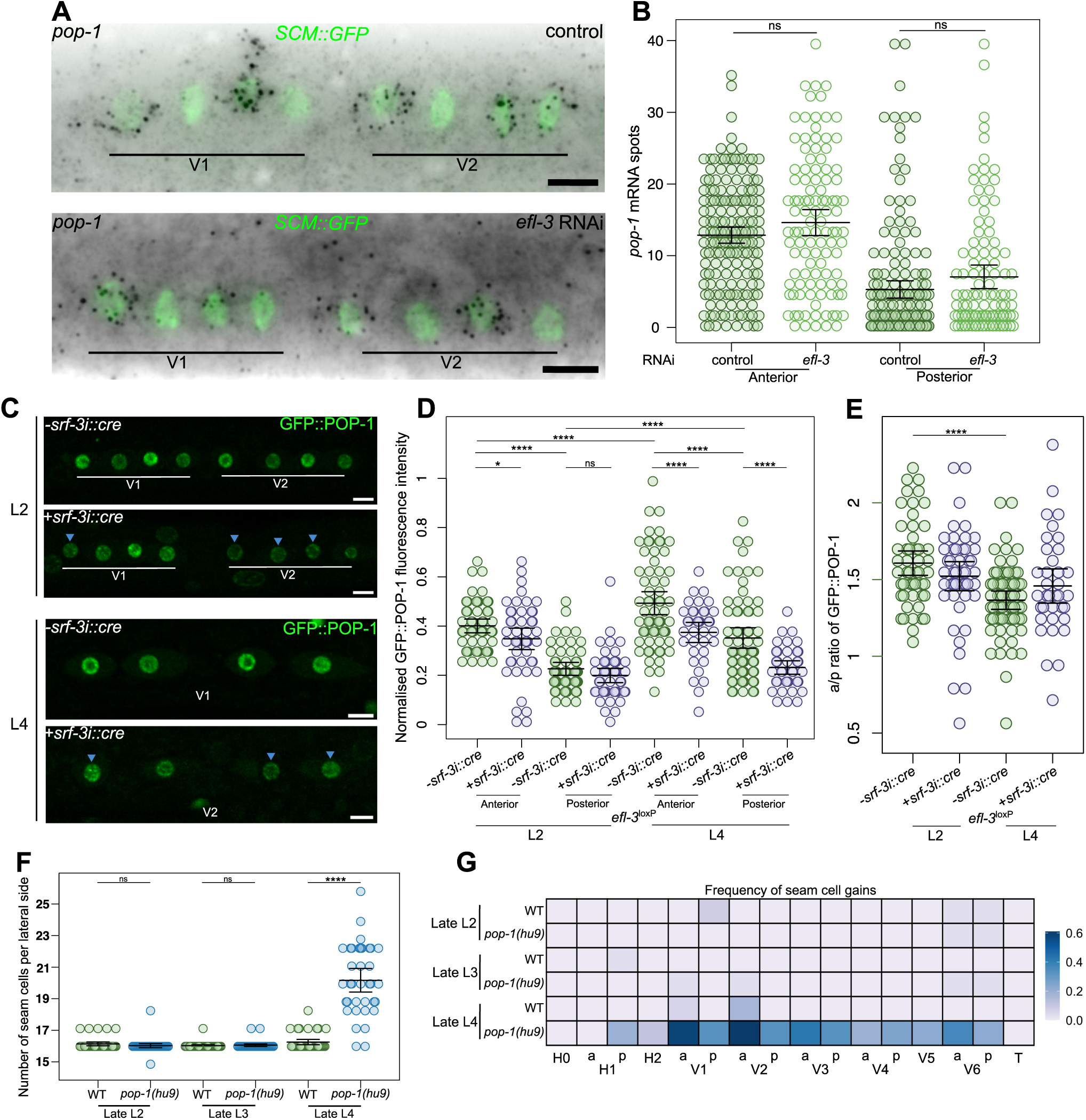
*efl-3* mutants show changes in POP-1 nuclear distribution at L2 and L4 stage. **(A)** Representative *pop-1* smFISH images after the L2 asymmetric division upon *efl-3* RNAi compared to control treatment. Seam cells are labelled using *SCM::GFP*. **(B)** Quantification of *pop-1* mRNA spots in V1-V4 lineages in anterior and posterior daughter cells following the L2 asymmetric division in the conditions of A, 108 ≤ n ≤ 174. **(C)** Representative images of GFP::POP-1 following the L2 and L4 asymmetric divisions in tissue-specific *efl-3* mutant animals (*+srf-3i::cre efl-3^loxP^*) versus wild-type (*-srf-3::cre efl-3^loxP^*). **(D)** Quantification of GFP::POP-1 fluorescence intensity following the L2 and L4 asymmetric divisions in tissue-specific *efl-3* mutant animals (*+srf-3i::cre efl-3^loxP^*) versus wild type (*-srf-3::cre efl-3^loxP^*), 36 ≤ n ≤ 65. * *p*<0.05 and **** *p*<0.001 with a t-test. **(E)** Anterior-posterior GFP::POP-1 ratios following the L2 and L4 asymmetric divisions in tissue-specific *efl-3* mutant animals versus wild type, 36 ≤ n ≤ 65. **** *p*<0.001 with a t-test. **(F)** Seam cell counts in wild-type and *pop-1(hu9)* mutant animals at the late L2, late L3 and late L4 stages. At late L4, *efl-3* mutants present a significant increase in the average number of seam cells compared to wild type, n = 36. **(G)** Heatmap showing frequency of seam cell gains per cell lineage at the late L2, late L3 and late L4 stages in wild-type and in *pop-1(hu9)* mutant animals., n = 36 per condition, a = anterior lineage, p = posterior lineage. Error bars in B, D, E and F show the mean ± standard deviation. Scale bars in A and C are 5 μm.

**Table S1: List of statistically significant EFL-3 NanoDam peaks.**

**Table S2: List of putative EFL-3 target genes in seam cells.**

**Table S3: List of smFISH probes used in this study.**

## STAR METHODS

### KEY RESOURCES TABLE

In an excel

### RESOURCE AVAILABILITY

#### Lead contact

Further information and requests for reagents should be directed to and will be fulfilled by the lead contact Michalis Barkoulas.

#### Materials availability

Plasmids and strains generated in this study are available upon request to the lead contact.

#### Data and code availability

### EXPERIMENTAL MODEL AND STUDY PARTICIPANT DETAILS

#### *C. elegans* maintenance and genetics

All *C. elegans* strains in this study were maintained at 20°C on Nematode Growth Medium (NGM) plates seeded with *E. coli* OP50^89^. A list of strains used in this study is provided in the Key Resources Table.

### METHOD DETAILS

#### Molecular Cloning and transient transgenesis

To construct seam cell CRE expression vector pHIN32, *C. elegans* codon-optimised woCre recombinase gene was amplified from pDK56 with oligos Wcre_F and Wcre_R from pDK56. The amplicon was inserted by Gibson Assembly into pDK70 containing *srf-3i* promoter. pHIN32 was injected into *efl-3* floxed animals at 5 ng/μl with 5 ng/μL *myo-2::dsRed* as co-injection marker and 90 ng/μl BJ36 as carrier DNA.

To overexpress *lit-1* using the last intron of *arf-5* (*arf-5i*), *lit-1^d^* fragment was amplified from PHX7906 cDNA using the oligos (pes10)_lit-1_Fw and (unc54)_lit-1_Rv. The amplicon was inserted by Gibson Assembly into pIR5 to create pMFM20. pMFM20 was injected into JR667 at 20 ng/μl with 5 ng/μL *myo-2::dsRed* as co-injection marker and 85 ng/μl BJ36 as carrier DNA.

#### RNAi feeding

RNAi by feeding was performed using *E. coli* HT115 expressing double-stranded RNA corresponding to the target gene or the control dsRNA plasmid (empty vector) as food source ^90^. Bacteria cultures were grown overnight and 300 μl were seeded onto NGM plates containing 25 μg/ml ampicillin, 12.5 μg/ml tetracyclin and 1 mM IPTG. All RNAi treatments were performed during the postembryonic development by seeding eggs from the corresponding strain onto the RNAi plates. RNAi bacteria clones used in this study are from the commercially available Ahringer RNAi Library^91^ (Source Bioscience).

#### Stable transgenesis and CRISPR genome editing

The endogenous copy of both *efl-3* and *lit-1* was tagged with codon optimized GFP and the auxin-inducible degradation (AID) domain just before the stop codon at the C-terminus using a CRISPR/Cas-9 approach (Suny Biotech).

*LoxP* sites flanking the *efl-3* endogenous locus (178 bp upstream of the transcription start site and 351 bp downstream of the 3’UTR) were introduced by injecting JR667 animals with 0.25 ng/μL protein Cas-9, 2.25 μM tracrRNA, 2.38 μM of 5’_efl-3_crRNA and 3’_efl-3_crRNAs, 5.5 μM of 5’ and 3’ repair templates and 5 ng/μL *myo-2::dsRed*. crRNAs targeted 241 bp upstream the TSS and 298 bp downstream the TES. 5’ and 3’ *efl-3* repair templates presented 50 bp homology to either side of the Cas-9 cut site and included a *loxP* sequence, a ClaI cut site and a M13uni_43 or M13rev_49 sequence, respectively. Transgenic animals with red pharynx were genotyped using the oligos efl-3_5p_F, R and efl-3_3p_F, R for 5’ and 3’ modification, respectively. The PCR product was digested with ClaI restriction enzyme to confirm successful homologous repair.

To delete the predicted EFL-3 binding region on *lit-1* locus, PHX7906 animals were injected with 0.25 ng/μL protein Cas-9, 2.25 μM tracrRNA, 2.38 μM of 5’_lit-1_crRNA and 3’_lit-1_crRNA and 5 ng/μL *myo-2::dsRed.* crRNAs targeted a 2743 bp long region inside *lit-1d* first intron. Transgenic animals with red pharynx were screened for the presence of the deletion using the oligos lit-1idel_Fw and lit-1idel_Rv. Deletion of the EFL-3 binding region was confirmed by sequencing.

Single-copy woCRE strain was generated with the MosSCI method using animals of the EG8080 strain with a Mos1 transposon insertion on chromosome III (*oxTi444*)^92^. pHIN32 was injected at 50 ng/μL together with 50 ng/μL of pCFJ601 (Mos1 transposase), 10 ng/μL of pMA122 (heat-shock-inducible *peel-1* toxin) and and 10 ng/μl, 2.5 ng/μl and 5 ng/μl of the co-injection markers (pGH8, *myo-2::dsRed*, *myo-3::mCherry*). After injection animals were kept at 25°C until they were starved. Heat shock was performed at 33°C for 3.5 hours and the plates were allowed to recover for 3 hours at room temperature before ‘reverse chunking’. The next day the top of the laws on each chunck was screened for normal roaming animals with the absence of co-injection markers. Homozygous lines were confirmed molecularly for the presence of single-copy transgenes using oligos NM3880 and NM3884.

#### Phenotypic analysis and microscopy

For phenotypic characterisation, animals were anesthetised on 3% agarose pads containing 100 μM sodium azide (NaN_3_), secured with a coverslip and visualised under an AxioScope A1 (Zeiss) upright epifluorescence microscope with a LED light source with a RETIGA R6 camera controlled by the Ocular software. Seam cell number was scored in 1 lateral side of early adult animals containing *SCM::GFP* marker. Visualisation of the seam cell membrane was achieved with *wrt-2::GFP::PH; wrt-2::GFP::H2B* markers and animals were scored at 30 hours post-bleaching for late L2 stage, at 40 hours post-bleaching for late L3 stage and 48 hours post-bleaching for late L4 stage.

Protein reporter images were acquired using a Leica SP8 Inverted confocal microscope with a x63/1.4 oil DIC objective controlled by LAS X software. Animals were mounted in 5% agarose pads containing 100 μM sodium azide (NaN_3_) and secured with a coverslip. To visualise L2 symmetric and asymmetric divisions animals were observed at 26 hours post-bleaching. To visualise L3 asymmetric divisions, animals were synchronised by egg laying over a period of 2 hours and observed after 35 hours. Fluorescence intensity for EFL-3::GFP, LIT-1::GFP, EGL-18::mNG and GFP::POP-1 was quantified using FIJI macros developed by the Imperial College FILM facility. Picture editing was performed using straightening tool and stitching macro from FIJI^93^.

#### Single molecule fluorescent in situ hybridisation

Animals were synchronised by bleaching and were fixed at the appropriate stage using 4% formaldehyde for 45 minutes and permeabilised with 70% ethanol. Samples were then incubated with a pool of 24-48 oligos labelled with Quasar 670 (Biosearch Technologies, Novato, CA) for 16 hours. Imaging was performed using the x100 objective of an epifluorescence Ti-eclipse microscope (Nikon) and DU-934 CCD-17291 camera (Andor Technology) operated via the Nikon NIS Elements Software. 23 z-stack slices of 0.8 μm were acquired for each animal. Spot quantification was performed using MATLAB (MathWorks) as previously described ^94^. Z-slices containing spots where max-projected to give a single image, inverted and merged to the GFP channel using ImageJ (NIH). A complete list of the smFISH probes used in this study is provided in Table S3.

For DAPI intensity quantification, z-slices were average-projected. DAPI fluorescence intensity was quantified by manually drawing a region of interest around the seam cell nuclei and measurements of average intensity were taken. Three background readings were taken from the area surrounding the seam cells.

#### Extraction and amplification of methylated DNA

For Nanodam experiments animals were synchronised by bleaching and grown for three generations in *E. coli* dam-/- dcm-/- bacteria (New England Biolabs, C2925). Two biological replicates were grown in parallel for strains carrying the NanoDam transgenes with and without *efl-3:gfp* for C-terminal and N-terminal Dam:GFP-nanobody configurations. Animals were collected after 48 hours, extensively washed x5 in M9 buffer and stored at −20°C. Purification and amplification of methylated DNA was performed as previously described^56^. Extracted gDNA was digested with DpnI and double stranded adaptors were ligated using T4 DNA ligase. Then, DNA fragments were digested with DpnII and amplified using MyTaq polymerase. Amplicons were purified using QIAquick PCR purification kit and adaptors were removed by AlwI digestion followed by another purification. Library preparation and Next Generation Sequencing on an Illumina HiSeq 4000 platform was performed by GENEWIZ.

#### NanoDam signal profiles, peak calling, and gene assignment

FASTQ files representing paired-end reads for each sample and replicate were processed using the damidseq_pipeline (v.1.5.3)^63^. Reads were mapped to the *C. elegans* WBcel235 genome assembly using bowtie2 (v.2.4.5) and scores were calculated per GATC fragment of the genome. Pairwise genome-wide log_2_(*efl-3-GFP*; *nanobody-dam*/ *nanobody-dam*) calculations were performed from bam files to produce bedGraph signal files that were normalised by quantile normalisation and arithmetically averaged into a single signal profile. Peak calling was performed using the perl script find_peaks (available at https://github.com/owenjm/find_peaks) with FDR < 0.05. Statistically significant peaks were assigned to genes using UROPA ^95^ with Caenorhabditis_elegans.WBcel235.106.gtf as the genome annotation file. Peaks were assigned to genes when their centre coordinate was positioned up to 5 kb upstream of a gene start site or 2 kb downstream of the end site. Visualisation of signal tracks and other genomic features was performed using Integrative Genomics Viewer (IGV).

#### Aggregation plots and heatmaps of signal localization

Signal aggregation plots and peak localisation heatmaps were generated using the SeqPlots GUI application^96^. Signal and peaks were visualised across genes containing peaks up to 5kb upstream of a gene start site or 2 kb downstream of the end site. Genes were plotted as “anchored” features, where the TSS and the TES are fixed in the x axis and the genic sequence is adjusted to a pseudo-length of 2kb. In aggregation plots, signal is averaged in 10bp bins and y axis represents deviations from the average signal across the plotted region (z-scores). In heatmaps, each coloured line represents the position of a statistically significant peak.

#### Assessment of overlap between statistically significant peaks

To calculate overlapping statistically significant peaks between the C-terminal and N-terminal Dam:GFP-nanobody configurations, BEDTools intersect tool was used^97^. To test whether overlaps where statistically significant, Monte Carlo simulations were performed using OLOGRAM^98^.

### QUANTIFICATION AND STATISTICAL ANALYSIS

Graphic representation and statistical analysis were performed using R. Error bars used in all graphs represent the standard deviation. Heatmaps represent relative frequency of seam cell defects per cell lineage, which was calculated by dividing the total number of gains or losses in a cell lineage by the total amount of animals analysed. An unpaired t-test or chi-square test was used to evaluate significance as indicated in the figure legends. Results were considered statistically significant when p < 0.05. Asterisks in figures indicate corresponding statistical significance as it follows: * p < 0.05; ** p < 0.01; *** p < 0.005; **** p < 0.001.

## ACKNOWLEDGEMENTS

This work was supported by the Wellcome Trust [219448/Z/19/Z] and a studentship by La Caixa Foundation. We thank Mark Hintze and Dimitris Katsanos for reagents and bioinformatic support. Some strains were provided by the CGC, which is funded by NIH Office of Research Infrastructure Programs (P40 OD010440). We thank Suny Biotech for generating the EFL-3::GFP and LIT-1::GFP strains. We also thank the facility for Imaging by Light Microscopy (FILM) at Imperial College London, which is part-supported by funding from the Wellcome Trust [104931/Z/14/Z] and the BBSRC [BB/T017929/1].

## Data and Materials Availability

All unique/stable reagents generated in this study are available from the Lead Contact without restriction.

The raw NanoDam sequence files and processed signal files have been deposited in the National Center for Biotechnology Information Gene Expression Omnibus under accession number GSE270959.

## Contact for Reagent and Resource Sharing

Further information and requests for resources and reagents should be directed to and will be fulfilled by the Lead Contact, Michalis Barkoulas.

## References

1. Slack, J.M.W. (2018). The science of stem cells (Blackwell) 10.1002/9781119235293.

2. Morrison, S.J., and Kimble, J. (2006). Asymmetric and symmetric stem-cell divisions in development and cancer. Nature 441, 1068–1074. 10.1038/nature04956.

3. Bajaj, J., Diaz, E., and Reya, T. (2020). Stem cells in cancer initiation and progression. Journal of Cell Biology 219, 1–12. 10.1083/jcb.201911053.

4. Joshi, P.M., Riddle, M.R., Djabrayan, N.J.V., and Rothman, J.H. (2010). *Caenorhabditis elegans* as a model for stem cell biology. Developmental Dynamics 239, 1539–1554. 10.1002/DVDY.22296.

5. Sulston, J.E., and Horvitz, H.R. (1977). Post-embryonic cell lineages of the nematode, *Caenorhabditis elegans*. Dev Biol 56, 110–156. 10.1016/0012-1606(77)90158-0.

6. Hedgecock, E.M., and White, J.G. (1985). Polyploid Tissues in the Nematode *Caenorhabditis elegans*. Dev Biol 107, 128–133.

7. Chisholm, A.D., and Hsiao, T.I. (2012). The *Caenorhabditis elegans* epidermis as a model skin. I: Development, patterning, and growth. Wiley Interdiscip Rev Dev Biol 1, 861–878. 10.1002/wdev.79.

8. Altun, Z.F., and Hall, D.H. (2024). Epithelial system, hypodermis. In WormAtlas.

9. Katsanos, D., Koneru, S.L., Mestek Boukhibar, L., Gritti, N., Ghose, R., Appleford, P.J., Doitsidou, M., Woollard, A., van Zon, J.S., Poole, R.J., et al. (2017). Stochastic loss and gain of symmetric divisions in the *C. elegans* epidermis perturbs robustness of stem cell number. PLoS Biol 15, 1–31. 10.1371/journal.pbio.2002429.

10. Pani, A.M., and Goldstein, B. (2018). Direct visualization of a native Wnt in vivo reveals that a long-range Wnt gradient forms by extracellular dispersal. Elife 7, 1– 22. 10.7554/eLife.38325.

11. Gleason, J.E., and Eisenmann, D.M. (2010). Wnt signaling controls the stem cell-like asymmetric division of the epithelial seam cells during *C. elegans* larval development. Dev Biol 348, 58–66. 10.1016/j.ydbio.2010.09.005.

12. Yamamoto, Y., Takeshita, H., and Sawa, H. (2011). Multiple Wnts redundantly control polarity orientation in *Caenorhabditis elegans* epithelial stem cells. PLoS Genet 7. 10.1371/journal.pgen.1002308.

13. Jackson, B.M., and Eisenmann, D.M. (2012). β-catenin-dependent Wnt signaling in *C. elegans*: Teaching an old dog a new trick. Cold Spring Harb Perspect Biol 4, 1–18. 10.1101/cshperspect.a007948.

14. Sawa, H., and Korswagen, H.C. (2013). Wnt signaling in *C. elegans*. WormBook, 1–30. 10.1895/wormbook.1.

15. Goldstein, B., Takeshita, H., Mizumoto, K., and Sawa, H. (2006). Wnt signals can function as positional cues in establishing cell polarity. Dev Cell 10, 391–396. 10.1016/j.devcel.2005.12.016.

16. Green, J.L., Inoue, T., and Sternberg, P.W. (2008). Opposing Wnt Pathways Orient Cell Polarity during Organogenesis. Cell 134, 646–656. 10.1016/j.cell.2008.06.026.

17. Mizumoto, K., and Sawa, H. (2007). Two βs or not two βs: regulation of asymmetric division by β-catenin. Trends Cell Biol 17, 465–473. 10.1016/J.TCB.2007.08.004.

18. Baldwin, A.T., and Phillips, B.T. (2014). The tumor suppressor APC differentially regulates multiple β-catenins through the function of axin and CKIα during *C. elegans* asymmetric stem cell divisions. J Cell Sci 127, 2771–2781. 10.1242/jcs.146514.

19. Mizumoto, K., and Sawa, H. (2007). Cortical β-Catenin and APC Regulate Asymmetric Nuclear β-Catenin Localization during Asymmetric Cell Division in *C. elegans*. Dev Cell 12, 287–299. 10.1016/j.devcel.2007.01.004.

20. Takeshita, H., and Sawa, H. (2005). Asymmetric cortical and nuclear localizations of WRM-1/β-catenin during asymmetric cell division in *C. elegans*. Genes Dev 19, 1743–1748. 10.1101/gad.1322805.

21. Sugioka, K., Mizumoto, K., and Sawa, H. (2011). Wnt regulates spindle asymmetry to generate asymmetric nuclear β-catenin in *C. elegans*. Cell 146, 942–954. 10.1016/j.cell.2011.07.043.

22. Yang, X.D., Karhadkar, T.R., Medina, J., Robertson, S.M., and Lin, R. (2015). β-Catenin-related protein WRM-1 is a multifunctional regulatory subunit of the LIT-1 MAPK complex. Proc Natl Acad Sci U S A 112, E137–E146. 10.1073/pnas.1416339112.

23. Shin, Yasuda, J., Rocheleau, C.E., Lin, R., Soto, M., Bei, Y., Davis, R.J., and Mello, C.C. (1999). MOM-4, a MAP Kinase Kinase Kinase-Related Protein, Activates WRM-1/LIT-1 Kinase to Transduce Anterior/Posterior Polarity Signals in *C. elegans*. Mol Cell 4, 275–280.

24. Lo, M.-C., Gay, F., Odom, R., Shi, Y., and Lin, R. (2004). Phosphorylation by the-Catenin/MAPK Complex Promotes 14-3-3-Mediated Nuclear Export of TCF/POP-1 in Signal-Responsive Cells in *C. elegans*. Cell 117, 95–106.

25. Meneghini, M.D., Ishitani, T., Carter, C.J., Hisamoto, N., Ninomiya-Tsuji, J., Thorpe, C.J., Hamill, D.R., Matsumoto, K., and Bowerman, B. (1999). MAP kinase and Wnt pathways converge to downregulate an HMG-domain repressor in *Caenorhabditis elegans*. Nature 399, 793–797.

26. Rocheleau, C.E., Yasuda, J., Shin, T.H., Lin, R., Sawa, H., Okano, H., Priess, J.R., Davis, R.J., and Mello, C.C. (1999). WRM-1 Activates the LIT-1 Protein Kinase to Transduce Anterior/Posterior Polarity Signals in *C. elegans*. Cell 97, 717–726.

27. Yang, X.D., Huang, S., Lo, M.C., Mizumoto, K., Sawa, H., Xu, W., Robertson, S., and Lin, R. (2011). Distinct and mutually inhibitory binding by two divergent β-catenins coordinates TCF levels and activity in *C. elegans*. Development 138, 4255–4265. 10.1242/dev.069054.

28. Shetty, P., Lo, M.C., Robertson, S.M., and Lin, R. (2005). *C. elegans* TCF protein, POP-1, converts from repressor to activator as a result of Wnt-induced lowering of nuclear levels. Dev Biol 285, 584–592. 10.1016/j.ydbio.2005.07.008.

29. Huang, S., Shetty, P., Robertson, S.M., and Lin, R. (2007). Binary cell fate specification during *C. elegans* embryogenesis driven by reiterated reciprocal asymmetry of TCF POP-1 and its coactivator β-catenin SYS-1. Development 134, 2685–2695. 10.1242/dev.008268.

30. Phillips, B.T., III, A.R.K., King, R., Hardin, J., and Kimble, J. (2007). Reciprocal asymmetry of SYS-1/β-catenin and POP-1/TCF controls asymmetric divisions in *Caenorhabditis elegans*. Proc Natl Acad Sci U S A 104, 3231–3236. 10.1073/pnas.0611507104.

31. Gorrepati, L., Thompson, K.W., and Eisenmann, D.M. (2013). *C. elegans* GATA factors EGL-18 and ELT-6 function downstream of Wnt signaling to maintain the progenitor fate during larval asymmetric divisions of the seam cells. Development (Cambridge) 140, 2093–2102. 10.1242/dev.091124.

32. Bekas, K.N., and Phillips, B.T. (2022). unc-37/Groucho and lsy-22/AES repress Wnt target genes in *C. elegans* asymmetric cell divisions. bioRxiv, 2022.01.10.475695. 10.1101/2022.01.10.475695.

33. Huang, X., Tian, E., Xu, Y., and Zhang, H. (2009). The *C. elegans* engrailed homolog *ceh-16* regulates the self-renewal expansion division of stem cell-like seam cells. Dev Biol 333, 337–347. 10.1016/j.ydbio.2009.07.005.

34. Kagoshima, H., Nimmo, R., Saad, N., Tanaka, J., Miwa, Y., Mitani, S., Kohara, Y., and Woollard, A. (2007). The *C. elegans* CBFβ homologue BRO-1 interacts with the Runx factor, RNT-1, to promote stem cell proliferation and self-renewal. Development 134, 3905–3915. 10.1242/dev.008276.

35. Katsanos, D., and Barkoulas, M. (2022). Targeted DamID in *C. elegans* reveals a direct role for LIN-22 and NHR-25 in antagonizing the epidermal stem cell fate. Sci Adv 8, 1–16. 10.1126/sciadv.abk3141.

36. Koh, K., and Rothman, J.H. (2001). ELT-5 and ELT-6 are required continuously to regulate epidermal seam cell differentiation and cell fusion in *C. elegans*. Development 128, 2867–2880.

37. Nimmo, R., and Woollard, A. (2008). Worming out the biology of Runx. Dev Biol 313, 492–500. 10.1016/j.ydbio.2007.11.002.

38. Smith, J.A., McGarr, P., and Gilleard, J.S. (2005). The *Caenorhabditis elegans* GATA factor elt-1 is essential for differentiation and maintenance of hypodermal seam cells and for normal locomotion. J Cell Sci 118, 5709–5719. 10.1242/jcs.02678.

39. Xia, D., Zhang, Y., Huang, X., Sun, Y., and Zhang, H. (2007). The *C. elegans* CBFβ homolog, BRO-1, regulates the proliferation, differentiation and specification of the stem cell-like seam cell lineages. Dev Biol 309, 259–272. 10.1016/j.ydbio.2007.07.020.

40. Attwooll, C., Denchi, E.L., and Helin, K. (2004). The E2F family: Specific functions and overlapping interests. EMBO Journal 23, 4709–4716. 10.1038/sj.emboj.7600481.

41. Van Den Heuvel, S., and Dyson, N.J. (2008). Conserved functions of the pRB and E2F families. Nat Rev Mol Cell Biol 9, 713–724. 10.1038/nrm2469.

42. Di Stefano, L., Jensen, M.R., and Helin, K. (2003). E2F7, a novel E2F featuring DP-independent repression of a subset of E2F-regulated genes. EMBO Journal 22, 6289–6298. 10.1093/emboj/cdg613.

43. Christensen, J., Cloos, P., Toftegaard, U., Klinkenberg, D., Bracken, A.P., Trinh, E., Heeran, M., Di Stefano, L., and Helin, K. (2005). Characterization of E2F8, a novel E2F-like cell-cycle regulated repressor of E2F-activated transcription. Nucleic Acids Res 33, 5458–5470. 10.1093/NAR/GKI855.

44. Logan, N., Graham, A., Zhao, X., Fisher, R., Maiti, B., Leone, G., and La Thangue, N.B. (2005). E2F-8: An E2F family member with a similar organization of DNA-binding domains to E2F-7. Oncogene 24, 5000–5004. 10.1038/sj.onc.1208703.

45. De Bruin, A., Maiti, B., Jakoi, L., Timmers, C., Buerki, R., and Leone, G. (2003). Identification and Characterization of E2F7, a Novel Mammalian E2F Family Member Capable of Blocking Cellular Proliferation. Journal of Biological Chemistry 278, 42041–42049. 10.1074/jbc.M308105200.

46. Lammens, T., Li, J., Leone, G., and De Veylder, L. (2009). Atypical E2Fs: new players in the E2F transcription factor family. Trends Cell Biol 19, 111–118. 10.1016/j.tcb.2009.01.002.

47. Kent, L.N., and Leone, G. (2019). The broken cycle: E2F dysfunction in cancer. Nat Rev Cancer 19, 326–338. 10.1038/s41568-019-0143-7.

48. Xie, D., Pei, Q., Li, J., Wan, X., and Ye, T. (2021). Emerging Role of E2F Family in Cancer Stem Cells. Front Oncol 11. 10.3389/fonc.2021.723137.

49. Cam, H., and Dynlancht, D. (2003). Emerging roles for E2F: Beyond the G1/S transition and DNA replication. Cancer Cell 3, 311–316.

50. Zalmas, L.P., Zhao, X., Graham, A.L., Fisher, R., Reilly, C., Coutts, A.S., and La Thangue, N.B. (2008). DNA-damage response control of E2F7 and E2F8. EMBO Rep 9, 252–259. 10.1038/sj.embor.7401158.

51. Chen, Z., and Han, M. (2001). *C. elegans* Rb NuRD and Ras regulate lin-39-mediated cell fusion during vulval fate specification. Current Biology 11, 1874– 1879. 10.1016/S0960-9822(01)00596-6.

52. Page, B.D., Guedes, S., Waring, D., and Priess, J.R. (2001). The *C. elegans* E2F- and DP-related proteins are required for embryonic asymmetry and negatively regulate Ras/MAPK signaling. Mol Cell 7, 451–460. 10.1016/S1097-2765(01)00193-9.

53. Ceol, C.J., and Horvitz, H.R. (2001). *dpl-1* DP and *efl-1* E2F act with *lin-35* Rb to antagonize Ras signaling in *C. elegans* vulval development. Mol Cell 7, 461–473. 10.1016/S1097-2765(01)00194-0.

54. Kudron, M., Niu, W., Lu, Z., Wang, G., Gerstein, M., Snyder, M., and Reinke, V. (2013). Tissue-specific direct targets of *Caenorhabditis elegans* Rb/E2F dictate distinct somatic and germline programs. Genome Biol 14, 1–17. 10.1186/gb-2013-14-1-r5.

55. Schertel, C., and Conradt, B. (2007). *C. elegans* orthologs of components of the RB tumor suppressor complex have distinct pro-apoptotic functions. Development 134, 3691–3701. 10.1242/dev.004606.

56. Katsanos, D., Ferrando-Marco, M., Razzaq, I., Aughey, G., Southall, T., and Barkoulas, M. (2021). Gene expression profiling of epidermal cell types in *C. elegans* using Targeted-DamID. Development 148, dev199452. 10.1242/DEV.199452.

57. Winn, J., Carter, M., Avery, L., and Cameron, S. (2011). Hox and a newly identified E2F co-repress cell death in *Caenorhabditis elegans*. Genetics 188, 897–905. 10.1534/genetics.111.128421.

58. Chen, H.Z., Ouseph, M.M., Li, J., Pécot, T., Chokshi, V., Kent, L., Bae, S., Byrne, M., Duran, C., Comstock, G., et al. (2012). Canonical and atypical E2Fs regulate the mammalian endocycle. Nat Cell Biol 14, 1192–1202. 10.1038/ncb2595.

59. Pandit, S.K., Westendorp, B., Nantasanti, S., Van Liere, E., Tooten, P.C.J., Cornelissen, P.W.A., Toussaint, M.J.M., Lamers, W.H., and De Bruin, A. (2012). E2F8 is essential for polyploidization in mammalian cells. Nat Cell Biol 14, 1181– 1191. 10.1038/ncb2585.

60. Yee, C., Xiao, Y., Katsanos, D., Medwig-Kinney, T.N., Zhang, W., Shen, K., Matus, D.Q., and Barkoulas, M. (2023). A NanoDam toolkit for tissue-specific transcription factor profiling in *C. elegans*. bioRxiv. 10.1101/2023.05.31.543105.

61. Tang, J.L.Y., Hakes, A.E., Krautz, R., Suzuki, T., Contreras, E.G., Fox, P.M., and Brand, A.H. (2022). NanoDam identifies novel temporal transcription factors conserved between the *Drosophila* central brain and visual system. Dev Cell 57, 1193–1207.

62. Southall, T.D., Gold, K.S., Egger, B., Davidson, C.M., Caygill, E.E., Marshall, O.J., and Brand, A.H. (2013). Cell-type-specific profiling of gene expression and chromatin binding without cell isolation: Assaying RNA pol II occupancy in neural stem cells. Dev Cell 26, 101–112. 10.1016/j.devcel.2013.05.020.

63. Marshall, O.J., and Brand, A.H. (2015). Damidseq-pipeline: An automated pipeline for processing DamID sequencing datasets. Bioinformatics 31, 3371–3373. 10.1093/bioinformatics/btv386.

64. Cao, J., Packer, J.S., Ramani, V., Cusanovich, D.A., Huynh, C., Daza, R., Qiu, X., Lee, C., Furlan, S.N., Steemers, F.J., et al. (2017). Comprehensive single-cell transcriptional profiling of a multicellular organism. Science (1979) 357, 661–667. 10.1126/science.aam8940.

65. Ashley, G.E., Duong, T., Levenson, M.T., Martinez, M.A.Q., Johnson, L.C., Hibshman, J.D., Saeger, H.N., Palmisano, N.J., Doonan, R., Martinez-Mendez, R., et al. (2021). An expanded auxin-inducible degron toolkit for *Caenorhabditis elegans*. Genetics 217. 10.1093/genetics/iyab006.

66. Coutts, A.S., Munro, S., and La Thangue, N.B. (2018). Functional interplay between E2F7 and ribosomal rRNA gene transcription regulates protein synthesis. Cell Death Dis 9. 10.1038/s41419-018-0529-6.

67. Zalmas, L.P., Coutts, A.S., Helleday, T., and Thangue, N.B.L. (2013). E2F-7 couples DNA damage-dependent transcription with the DNA repair process. Cell Cycle 12, 3037–3051. 10.4161/cc.26078.

68. Kipreos, E.T., and van den Heuvel, S. (2019). Developmental control of the cell cycle: Insights from *Caenorhabditis elegans*. Genetics 211, 797–829. 10.1534/genetics.118.301643.

69. Thurlings, I., Martínez-López, L.M., Westendorp, B., Zijp, M., Kuiper, R., Tooten, P., Kent, L.N., Leone, G., Vos, H.J., Burgering, B., et al. (2017). Synergistic functions of E2F7 and E2F8 are critical to suppress stress-induced skin cancer. Oncogene 36, 829–839. 10.1038/onc.2016.251.

70. Park, S.A., Lim, Y.J., Ku, W.L., Zhang, D., Cui, K., Tang, L.Y., Chia, C., Zanvit, P., Chen, Z., Jin, W., et al. (2022). Opposing functions of circadian protein DBP and atypical E2F family E2F8 in anti-tumor Th9 cell differentiation. Nat Commun 13. 10.1038/s41467-022-33733-8.

71. Segeren, H.A., van Rijnberk, L.M., Moreno, E., Riemers, F.M., van Liere, E.A., Yuan, R., Wubbolts, R., de Bruin, A., and Westendorp, B. (2020). Excessive E2F Transcription in Single Cancer Cells Precludes Transient Cell-Cycle Exit after DNA Damage. Cell Rep 33. 10.1016/j.celrep.2020.108449.

72. Westendorp, B., Mokry, M., Groot Koerkamp, M.J.A., Holstege, F.C.P., Cuppen, E., and De Bruin, A. (2012). E2F7 represses a network of oscillating cell cycle genes to control S-phase progression. Nucleic Acids Res 40, 3511–3523. 10.1093/nar/gkr1203.

73. Mitxelena, J., Apraiz, A., Vallejo-Rodriguez, J., Malumbres, M., and Zubiaga, A.M. (2016). E2F7 regulates transcription and maturation of multiple microRNAs to restrain cell proliferation. Nucleic Acids Res 44, 5557–5570. 10.1093/nar/gkw146.

74. Soukup, E.M., Bettinger, J.C., and Mathies, L.D. (2022). Transcription factors regulating the fate and developmental potential of a multipotent progenitor in *Caenorhabditis elegans*. G3: Genes, Genomes, Genetics 12. 10.1093/g3journal/jkac232.

75. van der Horst, S.E.M., Cravo, J., Woollard, A., Teapal, J., and van den Heuvel, S. (2019). *C. elegans* Runx/CBFβ suppresses POP-1 TCF to convert asymmetric to proliferative division of stem cell-like seam cells. Development 146, 1–14. 10.1242/dev.180034.

76. Maiti, B., Li, J., De Bruin, A., Gordon, F., Timmers, C., Opavsky, R., Patil, K., Tuttle, J., Cleghorn, W., and Leone, G. (2005). Cloning and characterization of mouse E2F8, a novel mammalian E2F family member capable of blocking cellular proliferation. J Biol Chem 280, 18211–18220. 10.1074/JBC.M501410200.

77. Van Rijnberk, L.M., Van Der Horst, S.E.M., Van Den Heuvel, S., and Ruijtenberg, S. (2017). A dual transcriptional reporter and CDK-activity sensor marks cell cycle entry and progression in *C. elegans*. PLoS One 12. 10.1371/journal.pone.0171600.

78. Jin, Y., Anbarchian, T., Wu, P., Sarkar, A., Fish, M., Peng, W.C., and Nusse, R. (2022). Wnt signaling regulates hepatocyte cell division by a transcriptional repressor cascade. PNAS 119. 10.1073/pnas.

79. Ishitani, T., Ninomiya-Tsuji, J., and Matsumoto, K. (2003). Regulation of Lymphoid Enhancer Factor 1/T-Cell Factor by Mitogen-Activated Protein Kinase-Related Nemo-Like Kinase-Dependent Phosphorylation in Wnt/β-Catenin Signaling. Mol Cell Biol 23, 1379–1389. 10.1128/mcb.23.4.1379-1389.2003.

80. Ishitani, T., Ninomiya-Tsuji, J., Nagai2, S.-I., Nishita2, M., Meneghini3, M., Barker, N., Watermank, M., Bowerman3, B., Clevers, H., Shibuya2, H., et al. (1999). The TAK1±NLK±MAPK-related pathway antagonizes signalling between β-catenin and transcription factor TCF. Nature 399, 798–802.

81. Ota, S., Ishitani, S., Shimizu, N., Matsumoto, K., Itoh, M., and Ishitani, T. (2012). NLK positively regulates Wnt/β-catenin signalling by phosphorylating LEF1 in neural progenitor cells. EMBO Journal 31, 1904–1915. 10.1038/emboj.2012.46.

82. Baldwin, A.T., Clemons, A.M., and Phillips, B.T. (2016). Unique and redundant β- catenin regulatory roles of two Dishevelled paralogs during *C. elegans* asymmetric cell division. J Cell Sci 129, 983–993. 10.1242/jcs.175802.

83. Ren, H., and Zhang, H. (2010). Wnt signaling controls temporal identities of seam cells in *Caenorhabditis elegans*. Dev Biol 345, 144–155. 10.1016/j.ydbio.2010.07.002.

84. Mallick, A., Ranawade, A., and Gupta, B.P. (2019). Role of PRY-1/Axin in heterochronic miRNA-mediated seam cell development. BMC Dev Biol 19, 1–12. 10.1186/s12861-019-0197-5.

85. Zhan, T., Rindtorff, N., and Boutros, M. (2017). Wnt signaling in cancer. Oncogene 36, 1461–1473. 10.1038/onc.2016.304.

86. Zhang, M., Xu, Y., Zhang, Y., and Lou, G. (2023). E2F8 knockdown suppresses cell proliferation and induces cell cycle arrest via Wnt/β-Catenin pathway in ovarian cancer. Chinese Journal of Physiology 66, 266–275. 10.4103/cjop.CJOP-D-22-00142.

87. Hughes, T.A., and Brady, H.J.M. (2005). Cross-talk between pRb/E2F and Wnt/β- catenin pathways: E2F1 induces axin2 leading to repression of Wnt signalling and to increased cell death. Exp Cell Res 303, 32–46. 10.1016/j.yexcr.2004.09.014.

88. Yu, S., Yerges-Armstrong, L.M., Chu, Y., Zmuda, J.M., and Zhang, Y. (2013). E2F1 effects on osteoblast differentiation and mineralization are mediated through up-regulation of frizzled-1. Bone 56, 234–241. 10.1016/j.bone.2013.06.019.

89. Brenner, S. (1974). The genetics of *Caenorhabditis elegans*. Genetics 77, 71–94.

90. Kamath, R.S., Fraser, A.G., Dong, Y., Poulin, G., Durbin, R., Gotta, M., Kanapin, A., Le Bot, N., Moreno, S., Sohrmann, M., et al. (2003). Systematic functional analysis of the *Caenorhabditis elegans* genome using RNAi. Nature 421, 231–237. 10.1038/nature01278.

91. Kamath, R.S., and Ahringer, J. (2003). Genome-wide RNAi screening in *Caenorhabditis elegans*. Methods 30, 313–321. 10.1016/S1046-2023(03)00050-1.

92. Frøkjær-Jensen, C., Davis, M.W., Sarov, M., Taylor, J., Flibotte, S., LaBella, M., Pozniakovsky, A., Moerman, D.G., and Jorgensen, E.M. (2014). Random and targeted transgene insertion in *Caenorhabditis elegans* using a modified Mos1 transposon. Nat Methods 11, 529–534. 10.1038/nmeth.2889.

93. Preibisch, S., Saalfeld, S., and Tomancak, P. (2009). Globally optimal stitching of tiled 3D microscopic image acquisitions. Bioinformatics 25, 1463–1465. 10.1093/bioinformatics/btp184.

94. Raj, A., van den Bogaard, P., Rifkin, S.A., van Oudenaarden, A., and Tyagi, S. (2008). Imaging individual mRNA molecules using multiple singly labelled probes. Nat Methods 5, 877–879. 10.1038/nmeth.1253.

95. Kondili, M., Fust, A., Preussner, J., Kuenne, C., Braun, T., and Looso, M. (2017). UROPA: A tool for Universal RObust Peak Annotation. Sci Rep 7. 10.1038/s41598-017-02464-y.

96. Stempor, P., and Ahringer, J. (2016). SeqPlots - Interactive software for exploratory data analyses, pattern discovery and visualization in genomics. Wellcome Open Res 1. 10.12688/WELLCOMEOPENRES.10004.1.

97. Quinlan, A.R., and Hall, I.M. (2010). BEDTools: a flexible suite of utilities for comparing genomic features. Bioinformatics 26, 841–842. 10.1093/BIOINFORMATICS/BTQ033.

98. Ferré, Q., Charbonnier, G., Sadouni, N., Lopez, F., Kermezli, Y., Spicuglia, S., Capponi, C., Ghattas, B., and Puthier, D. (2019). OLOGRAM: Determining significance of total overlap length between genomic regions sets. Bioinformatics 36, 1920–1922. 10.1093/BIOINFORMATICS/BTZ810.

